# Progressive alignment with Cactus: a multiple-genome aligner for the thousand-genome era

**DOI:** 10.1101/730531

**Authors:** Joel Armstrong, Glenn Hickey, Mark Diekhans, Alden Deran, Qi Fang, Duo Xie, Shaohong Feng, Josefin Stiller, Diane Genereux, Jeremy Johnson, Voichita Dana Marinescu, David Haussler, Jessica Alföldi, Kerstin Lindblad-Toh, Elinor Karlsson, Erich D. Jarvis, Guojie Zhang, Benedict Paten

## Abstract

Cactus, a reference-free multiple genome alignment program, has been shown to be highly accurate, but the existing implementation scales poorly with increasing numbers of genomes, and struggles in regions of highly duplicated sequence. We describe progressive extensions to Cactus that enable reference-free alignment of tens to thousands of large vertebrate genomes while maintaining high alignment quality. We show that Cactus is capable of scaling to hundreds of genomes and beyond by describing results from an alignment of over 600 amniote genomes, which is to our knowledge the largest multiple vertebrate genome alignment yet created. Further, we show improvements in orthology resolution leading to downstream improvements in annotation.

## 1 Introduction

New genome assemblies have been arriving at a rapidly increasing pace, thanks to rapid decreases in sequencing costs and improvements in third-generation sequencing technologies [8, 40, 22]. For example, the number of vertebrate genome assemblies currently in the NCBI database [25] has increased by over 50% in just the past year (to 1485 assemblies as of July 2019). The Vertebrate Genome Project, Genome 10K [26], the Earth BioGenome Project [31], the Bird 10K project [41], and the 200 Mammals project [14], among others, aim to release hundreds of high-quality assemblies of previously unsequenced genomes in the next year, and thousands over the next decade.

In addition to this influx of assemblies from different species, new human *de novo* assemblies [21] are being produced, which enable analysis of not just small polymorphisms, but also complex, large-scale structural differences between human individuals and haplotypes. This coming era and its unprecedented amount of data offers the opportunity to unlock many insights into genome evolution, but also presents challenges in adapting our analysis methods to meet the increased scale.

Often we want to make use of these assemblies to conduct analyses like species-tree inference, comparative annotation [11, 30], or constraint detection [19, 12]. All of these require comparing an assembly against one or more other assemblies. This involves creating a mapping from each region of each genome to a corresponding region in each other genome, taking into account the possibility of complex rearrangements: this is the problem of creating a *genome alignment* [1].

Genome aligners are one of the most fundamental tools used in comparative genomics, but since the problem is difficult, different aligners frequently give somewhat different results [6], and many intentionally limit the alignments they produce to simplify the problem. Two of the most common limitations are *reference-bias*, which constrains a multiple alignment to only regions present in a single reference genome, and restricting the alignment to be *single-copy*, which allows only a single alignment in any column in any given genome, causing the alignment to miss multiple-orthology relationships created by lineage-specific duplications. Cactus [35] is a genome alignment program which has neither of these restrictions; it is capable of generating a reference-free multiple alignment that allows detecting multiple-orthology relationships.

The version of Cactus available at the time performed very well in the Alignathon [6], an evaluation of genome aligners. However, the runtime of that initial iteration of Cactus scaled quadratically with the total number of bases in the alignment problem, making alignment of more than about ten vertebrate genomes completely impractical. To address these difficulties, we present fundamental changes to the Cactus process that incorporate a progressive alignment [10] strategy, which changes the runtime of the alignment to scale linearly with the number of genomes. We show that the result is an aligner that remains state-of-the-art in accuracy, and continues to lack reference bias, but which is tractable to use on hundreds to thousands of large, vertebrate-sized input genomes. This new version of Cactus has been developed over several years, and has already been successfully used as an integral component of high-profile comparative genomics projects [16, 5, 15, 32, 28]. We describe the many improvements to the original Cactus method that make large alignments tractable, while also increasing the accuracy of those alignments. We demonstrate it is capable of creating useful alignments across a wide range of evolutionary distances, from intra-species alignments useful in population genetics [13] to inter-species alignments spanning hundreds of millions of years of genome evolution. Because of its support for multiple-orthology relationships, it automatically supports diploid assemblies, which are becoming more common as new technologies enable phasing across long distances [27, 40].

## 2 Results

### 2.1 Cactus

The new progressive Cactus pipeline is freely available and open source. The only inputs needed are a guide tree and a FASTA file for each genome assembly.

The key innovation of the new Cactus aligner is to adapt the classic *progressive* strategy (used in collinear multiple alignment for decades [10]) to a whole-genome alignment setting. Progressive aligners use a *guide tree* to recursively break a multiple alignment problem into many smaller sub-alignments, each of which is solved independently; the resulting sub-alignments are themselves aligned together according to the tree structure to create the final alignment. Progressive alignment has been succesfully applied to whole-genome alignment before, for example by progressiveMauve [4] and TBA/MULTIZ [2]. Cactus now follows a similar strategy, with the key innovation being that Cactus implements a progressive-alignment strategy for whole-genome alignment using reconstructed ancestral assemblies as the method for combining sub-alignments. This strategy not only results in a much faster alignment runtime, but also produces ancestral reconstructions.

As a practical matter, Cactus also now uses the Toil [39] workflow framework to organize and distribute its computational tasks. Because it runs on Toil and supports container execution via Docker and Singularity [29], Cactus can be run on many different environments: single machines (for small alignments), conventional HPC clusters, as well as the GCP, AWS, and Azure clouds.

Figure 1A shows the overall organization of the new Cactus process. The guide tree, which need not be fully resolved (binary), is used to recursively split a large alignment problem (comparing every genome to every other genome) into many small subproblems, each of which compares only a small number (usually 2–5) of genomes against one another. The purpose of each subproblem is to reconstruct an ancestral assembly at each internal node in the guide tree, as well as to generate alignments between that internal node’s children and its ancestral reconstruction. The ancestral assemblies are then used as input genomes in subproblems further up the tree, while the parent-child alignments are later combined to produce the full alignment. Two sets of genomes are considered: the children of the internal node (which we call the *ingroup genomes*), and a set of non-descendants of that node (the *outgroup genomes*). The ingroup genomes form the core alignment relationship being established at this node. The outgroup genomes serve to answer the question of what sequence from the ingroups is also present in the ancestor (whether an indel among the ingroups is likely a deletion rather than an insertion), and in how many copies (whether a duplication predates or postdates the speciation event the node represents). The outgroups also provide information for guiding the ancestral assembly by providing additional order-and-orientation information, as well as additional information when generating ancestral base calls. These genome sets are used as the input to the main subproblem workflow, which we outline below and in Figure 1B, and describe in detail in Section 7.1.

**Figure 1:**
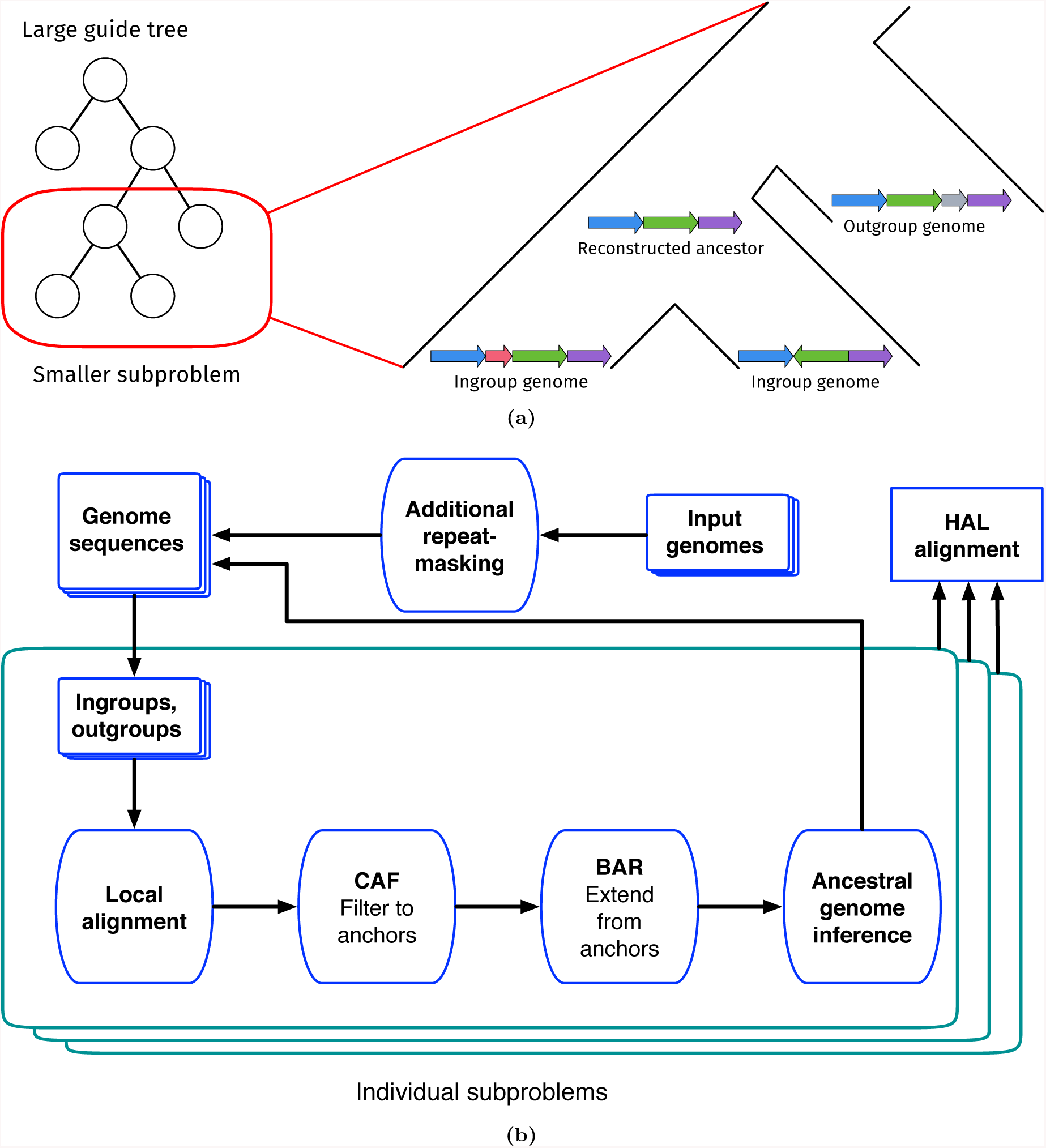
A diagram of the progressive process within Cactus. A: A large alignment problem is decomposed into many smaller subproblems using an input guide tree. Each subproblem compares a set of ingroup genomes (the children of the internal node to be reconstructed) against each other as well as a sample of outgroup genomes (non-descendants of the internal node in question). B: This flowchart represents the phases which the overall alignment, as well as each subproblem alignment, proceeds through. The end result is a new genome assembly representing Cactus’s reconstruction of the ancestral genome, as well as an alignment between this ancestral genome and its children. After all subproblems have been completed, the parent-child alignments are combined to create the full reference-free alignment in the HAL [18] format.

Each individual subproblem follows a procedure akin to the original Cactus process. The subproblem procedure begins with a set of pairwise local alignments generated via LASTZ [17]. These pairwise alignments are then filtered and combined into a cactus graph representing an initial multiple alignment using the CAF algorithm described in our earlier work [35], though we note important changes to the filtering in Section 7.1.4 and Section 7.1.5. The initial alignment is refined using the BAR algorithm again described in earlier work [35] to create a more complete alignment. The ancestral assembly is then created by ordering the blocks in this final alignment and establishing a most-likely base call for each column in each block. The resulting ancestral sequence is then fed into later subproblems (unless the subproblem represents the root of the guide tree, which indicates the end of the alignment process).

### 2.2 Evaluation on simulated data

To evaluate the improvements in quality and runtime of the alignments produced using the new progressive alignment strategy, we simulated the evolution of 20 30-megabase genomes using Evolver [7] along a tree of catarrhines. We ran two alignment strategies — one using a fully-resolved binary guide tree (which takes full advantage of the new progressive mode) and one using a fully-unresolved star guide tree (which is similar to the originally published version of Cactus) — across variously sized subsets of these genomes (for details of the simulation and alignments, see Section S1.1). The alignments using the progressive strategy were much faster, especially when aligning large numbers of genomes, as expected given its linear scaling runtime, as opposed to the quadratic scaling of the star-tree (Figure 2A). The simulated genomes have a known true alignment relating them, which is produced during the simulation process; using this it is possible to evaluate the quality of the alignments produced by the two strategies (Figure 2B). The progressive strategy is much more sensitive (89% recall) than the star strategy (82% recall) when aligning all 20 genomes: this reflects the fact that the increasing number of genomes will decrease the length of rearrangement-free regions, limiting the effectiveness of the rearrangement-based alignment filtering method that Cactus utilizes. Since the progressive strategy only aligns a constant number of genomes together at a time, it is able not only to make the runtime of aligning large numbers of genomes practical, but to align them with an accuracy unattainable by the previous versions of Cactus.

**Figure 2:**
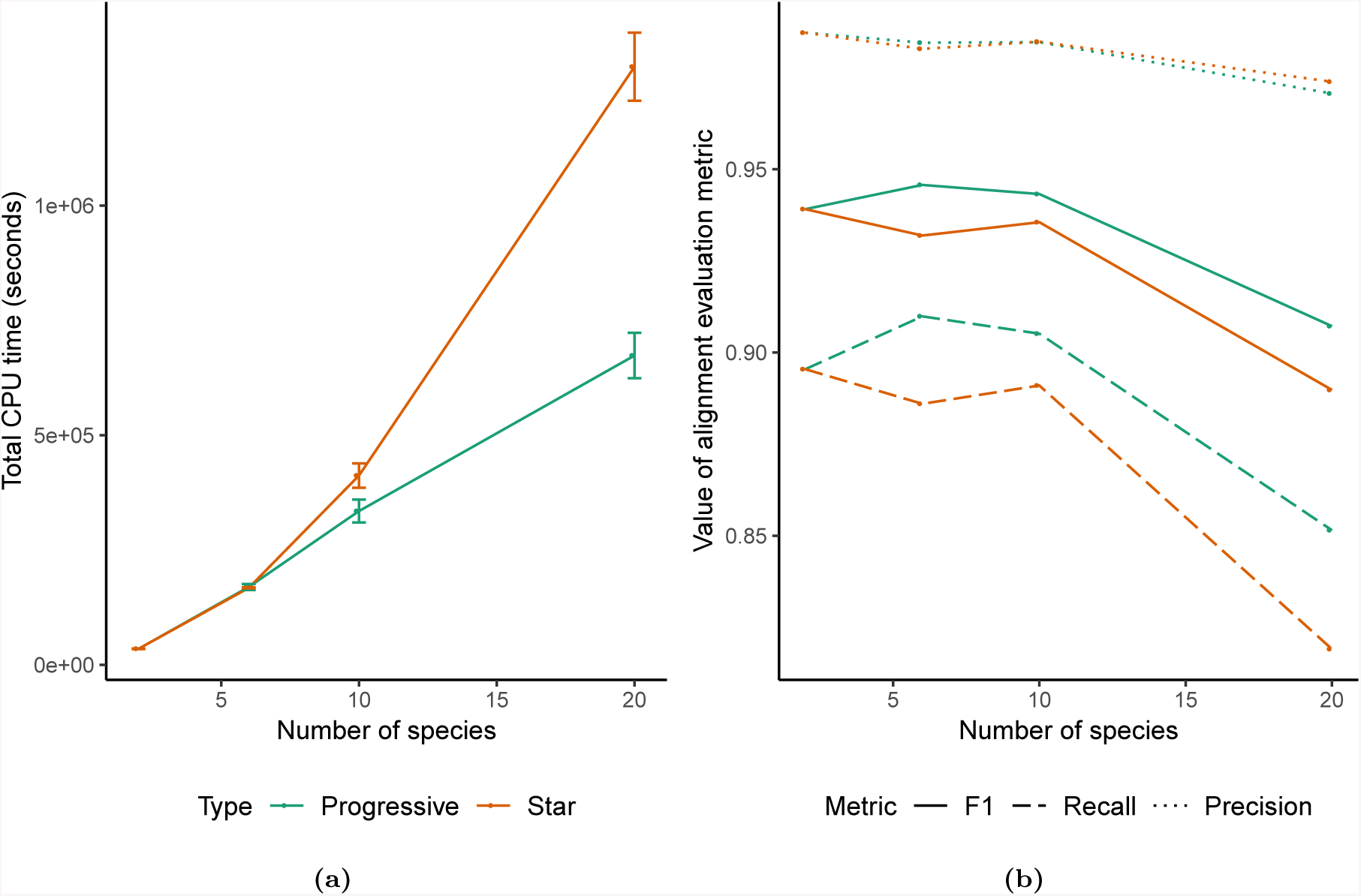
Results from alignments of varying numbers of simulated genomes using the progressive mode of Cactus (“Progressive”), versus the mode without progressive decomposition similar to originally described in [35] (“Star”). A) The total runtime of the two alignment methods across 3 runs. The runtime is nearly identical when aligning two genomes since the alignment problem is not further decomposed, but the linear scaling of the progressive mode means it is much faster with large numbers of genomes than the quadratic scaling required without progressive alignment. B) The precision, recall, and F1 score (harmonic mean of precision and recall) of aligned pairs for each alignment compared to pairs from the true alignment produced by the simulation.

### 2.3 Adding new genomes to an existing alignment

Given the rate of arrival of new assembly versions and newly sequenced genomes, adding new information to an alignment without recomputing it from scratch is valuable, especially for large alignments where recomputing the entire alignment is often cost-prohibitive. Cactus supports adding a new genome to an existing alignment by taking advantage of the tree structure of the progressive alignments it produces. There are three ways that a new genome can be added to an alignment, depending on its phylogenetic position relative to the existing genomes: 1) as outgroup to all the existing genomes in the alignment, 2) by being added as a new child of an existing ancestral genome in the alignment, or 3) by splitting a branch in the existing alignment, creating a new internal node and two new branches (Figure S1). Cactus allows adding a new genome in any of these ways, though the details differ; see Section 7.2. Assemblies can be replaced with new versions by simply deleting them and adding the new assembly in as a leaf. Adding multiple genomes is possible, either iteratively or (if the new genomes are monophyletic) by aligning together the new genomes and adding in the ancestral clade root.

We tested the effect of adding a new genome to an existing alignment using the same set of simulated catarrhine genomes as in Section 2.2. To replicate the use-case of an end-user wanting to add a genome to a previously-created alignment, we generated an alignment holding out one of the 20 genomes (the crab-eating macaque), and added that genome back into the alignment by both splitting an existing branch (resulting in the same topology as a full alignment would), and by adding the macaque as a new child of an existing ancestor (creating a trifurcation which did not exist in the original tree). For details of this process, see Section S1.2. Both methods resulted in alignments that had accuracy nearly identical to the full alignment that included the macaque from the start: both addition methods as well as the full alignment achieved an F1 score of 0.926 (Table 1).

**Table 1:** Results of adding a new genome to an alignment of simulated genomes. Precision, recall, and F1-score statistics are all of aligned pairs that contain a base of the added genome. An alignment where the genome was included initially is shown for comparison.

| Alignment | Precision | Recall | F1-score | CPU time |
| --- | --- | --- | --- | --- |
| Genome added to branch | 97.27% | 88.40% | 92.63% | 11 h |
| Genome added to node | 97.21% | 88.35% | 92.57% | 16 h |
| Full realignment of entire tree & new genome | 97.19% | 88.33% | 92.55% | 176 h |

### 2.4 Effect of the guide tree

Since Cactus uses an input guide tree to decompose the alignment problem, the guide tree can potentially impact the resulting alignment. This could be problematic when the exact species tree relating the input set of genomes is unknown or controversial. However, Cactus aims to reduce any effect of the guide tree by including a great deal of outgroup information, including multiple outgroups when possible. To quantify the effect of the guide tree on a large alignment with an uncertain species tree, we created four alignments of a set of 48 avian species, which we subsetted down to a single chromosome (Chromosome 1). The avian species tree is somewhat debated, with many different plausible hypotheses [23, 37], making birds an excellent test case with no single clearly correct guide tree. We aligned these birds using four different guide trees: two trees that represent two different hypotheses about the avian species tree [23, 37], one consensus tree between the former two trees, and one tree that was randomly permuted from the Jarvis et al. tree [23] (see Section S1.3 for details on the alignments and Figure S2 for a visualization of the four guide trees). The four alignments were highly similar, with an average of 98.5% of aligned pairs exactly identical between any two different alignments: detailed results are shown in Table 2.

**Table 2:** Comparison of alignment similarity between four alignments of the same 48 avian genomes with different guide trees. Similarity between each pair of alignments is represented by the F1 score (harmonic mean of precision and recall) of aligned-pair relationships in the two alignments.

| Guide tree | Jarvis | Prum | Consensus | Permuted |
| --- | --- | --- | --- | --- |
| Jarvis | 1 |  |  |  |
| Prum | 0.9867 | 1 |  |  |
| Consensus | 0.9882 | 0.9883 | 1 |  |
| Permuted | 0.9843 | 0.9822 | 0.9836 | 1 |

### 2.5 Timing duplication events

Users of a genome alignment are almost always interested in *orthology*, rather than *homology*, between a set of sequences. For example, when comparing human and chimpanzee KZNF genes, providing an alignment from each gene to the over-400 [20] homologous KZNF genes in the other genome is nigh-useless; the user is likely interested in only the orthologous copy or copies (in the case of a lineage-specific duplication) in the other genome. For this reason, Cactus alignments are capable of representing complex orthology/paralogy relationships, with an ability to display the alignment(s) labeled as orthologous, but also the option for a user to request alignments to paralogs at a customizable coalescence-time threshold. This is achieved by implicitly producing a gene tree as the alignment is built, albeit with some restrictions imposed by the output HAL [18] format, namely that a duplication event is represented by multiple regions in the child(ren) aligned to a single region in the parent species. This forbids the representation of gene-tree-species-tree discordance as would occur in incomplete lineage-sorting or horizontal transfer, as well as the exact ordering of multiple duplication events along a single branch. The restricted problem we solve at each subproblem step is that each block should represent all regions orthologous to a single region of the ancestral sequence, possibly multiple per species; we make no attempt to fully resolve the gene tree when multiple duplications take place along a single branch. However, this still requires resolving the timing of all duplication events: duplicated sequences whose coalescence precedes the speciation event represented in the subproblem should be split, while those following the speciation event should be kept together.

Because it is impractical to generate maximum-likelihood trees for every block in the subproblem, Cactus relies on heuristically filtering alignments to remove paralogs before building its cactus graph. For this we developed two heuristics: a filter based on similarity to outgroup sequence, which was used in the many projects which used the beta versions of progressive Cactus, and (more recently) a method of pre-filtering alignments that only allows any given base to contribute one “best” alignment in most cases (described in Section 7.1.4). Of the two methods, the newer best-hit filtering removes many more likely-paralogous alignments, especially to closely-related genomes, while leaving approximately the same amount of sequence covered by a single homology. For example, in two comparison alignments of the same 12 genomes, one using the best-hit filtering and one using the outgroup filtering, the amount of human sequence mapping to two or more places in the chimpanzee genome was reduced from 6.1% to 2.6%, while the total amount of human covered by chimpanzee actually increased despite the removed homologies (see Figure 3A,B for an example visualization and Figure 3C for aggregate statistics; see Section S1.4.1 for details on the alignments).

**Figure 3:**
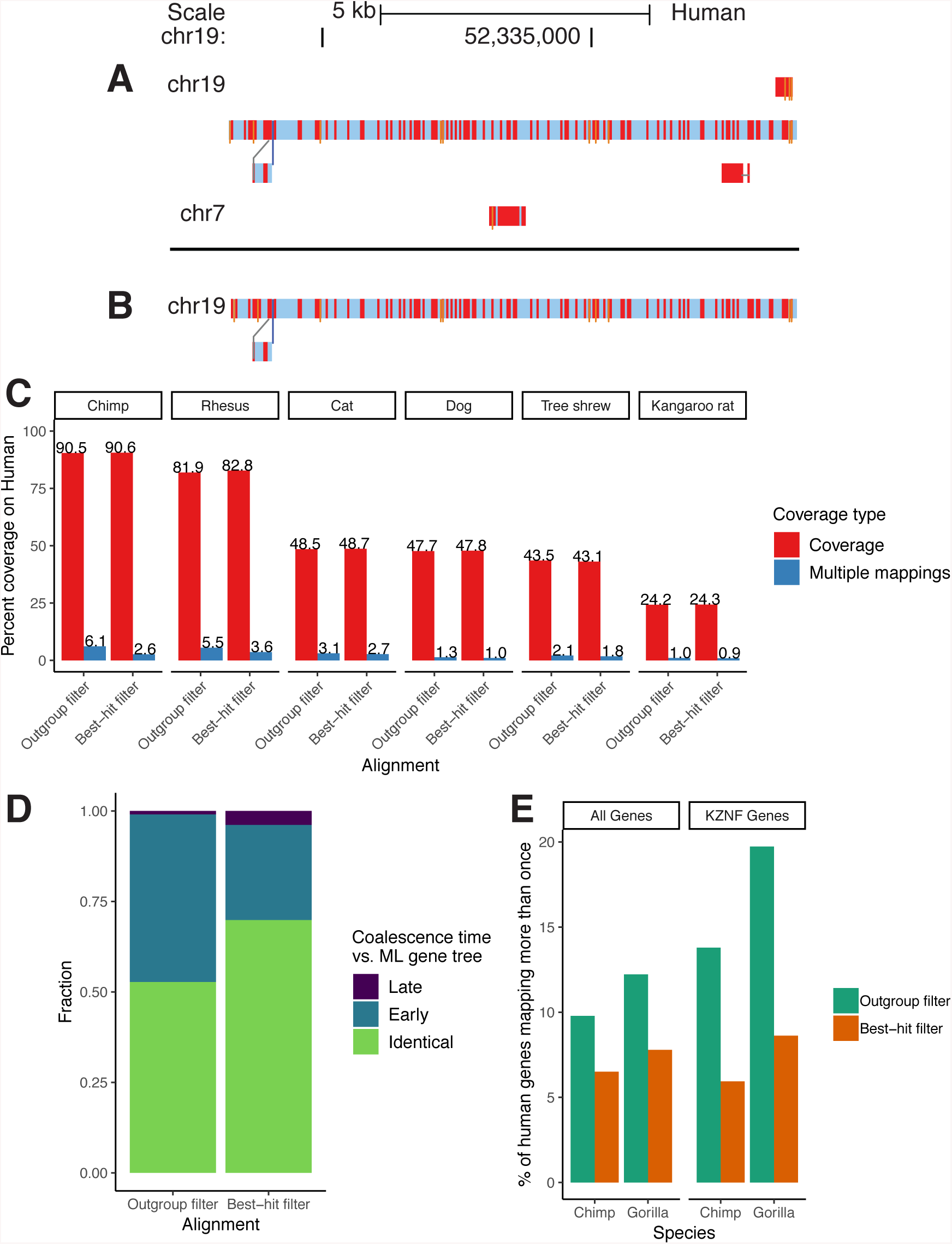
Results from the improved paralog-filtering method. A/B: A sample snake track [33] within a recently duplicated region before (A) and after (B) the filtering change. Nucleotide subsitutions are shown as red bars, and insertions are shown as thin orange bars. C: Coverage results from two alignments of identical assemblies using the outgroup and best-hit filtering methods. Multiple-mappings: sites which map to two or more sites on the target genome. D: Results from comparing phylogenetic trees implicit in the HAL alignment to ML re-estimated trees of the same regions. “Early” coalescences imply that too many duplication events have been created in the reconciled trees, while “Late” implies that too many loss events have been created. E: Percent of human genes that map more than once to the chimp/gorilla genomes in two CAT [11] annotations using alignments created with the outgroup and best-hit filtering methods. KZNF: KRAB zinc-finger genes.

To confirm that these improvements were likely caused by removal of paralogous rather than orthologous alignments, we compared phylogenetic trees implicit in the columns of HAL alignments to independently re-estimated approximately-ML trees produced by FastTree [36] for the same regions Section S1.4.3. Since HAL does not produce a fully binarized history of duplication events, we compared the species assigned to the most recent common ancestor (MRCA) of randomly selected pairs of sites from genomes containing a duplication within the column. If the species assigned to the MRCA in the HAL tree is a descendant of the species within the reconciled ML tree, that implies that there are paralogs represented as orthologs within the HAL tree (since a duplication event must have been resolved too early). Similarly, if the MRCA species within the HAL tree is an ancestor of that within the reconciled ML tree, a duplication event must have been resolved too late in the HAL, implying additional false loss / deletion events. The number of paralogous alignments (represented by the coalescence time between duplicated sequences being too “early” in the HAL tree relative to the ML tree) in the alignment of the 12 boreoeutherian genomes was clearly reduced (46% in the outgroup filtering vs 26% in the best-hit filtering) (Figure 3D).

We separately ran the Comparative Annotation Toolkit (CAT) [11] on identical chimpanzee and gorilla assemblies in two alignments using the outgroup and best-hit filtering methods (Section S1.4.2). Not only was CAT less likely to identify a human gene in multiple chimp loci using the best-hit filtering (e.g. 6.5% vs. 9.8% multiple-mappings across all genes in chimp, and 5.9% vs. 13.8% for the recently-duplicated KRAB zinc-finger gene family) (Figure 3E), but as a result orthologs for 104 more human genes were identified in the output gene set for chimp (182 in gorilla) (Table S3). This is likely because tens of thousands fewer paralogous transcripts were filtered out in the initial filtering phase of CAT (Table S2), reducing confusion about which transcript projection to put into the gene set.

### 2.6 600-way amniote alignment

To demonstrate this new version of Cactus we present results from an alignment of 605 amniote genomes, relating in a reference-free manner a total of over 1 trillion bases of DNA across hundreds of millions of years of genome evolution. The amniote-wide alignment combines two smaller alignments: one created for the 200 Mammals project [14], representing 242 placental mammals, and one for the Bird 10K project [41], which relates 363 avians. The overall topology is shown in Figure 4A. To our knowledge this represents the largest whole-genome alignment yet created. Table 3 contains aggregate statistics on this alignment, which was computed using the Amazon Web Services (AWS) cloud infrastructure (for details on the construction, see Section S1.6).

**Figure 4:**
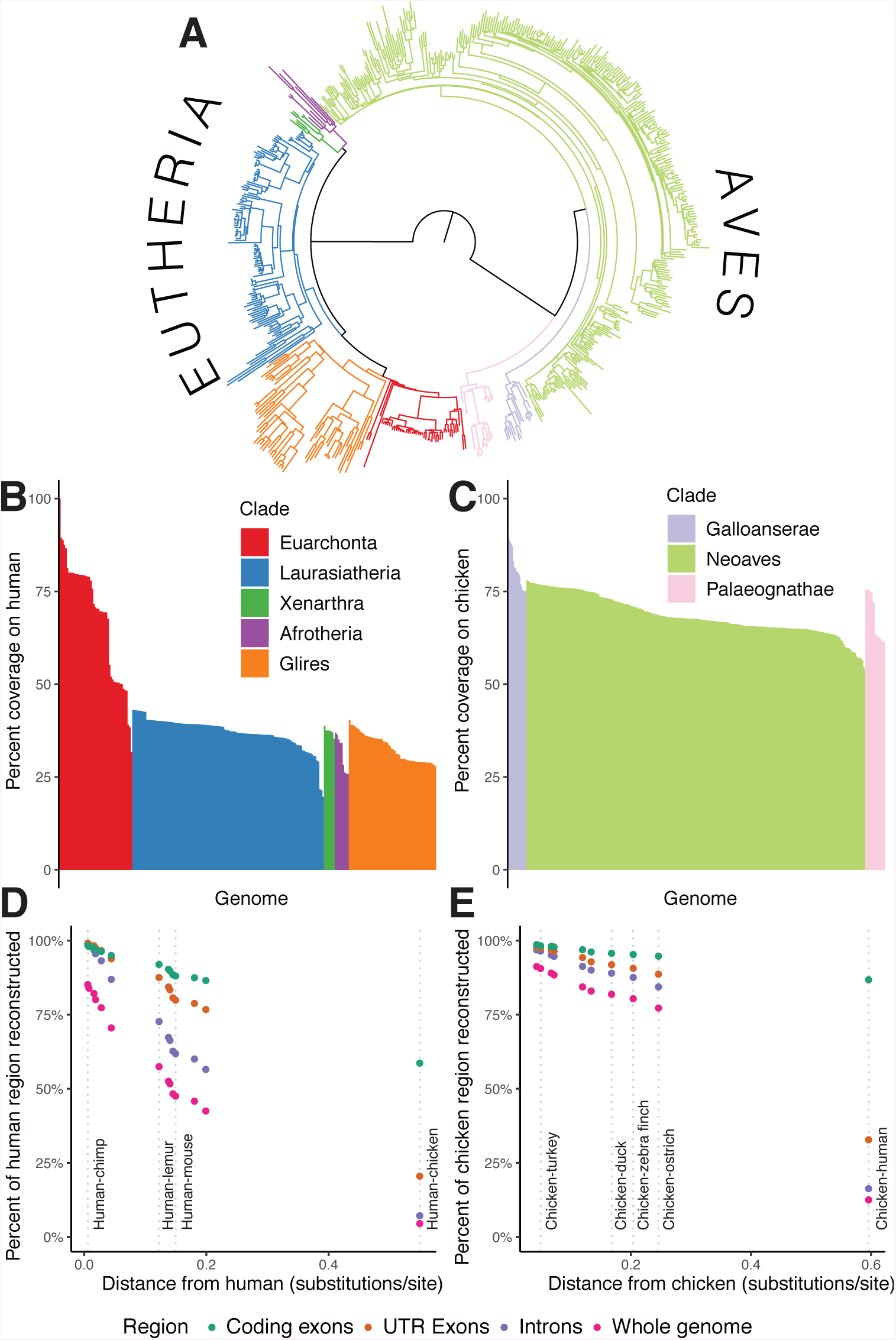
Results from the 600-way amniote alignment. A: The species tree relating the 600 genomes. Branches are colored by clades in the same way as figures B and C. B: Percent coverage on human within the eutherian mammals, grouped by clade from highest to lowest coverage. C: Similar to B, but for coverage on chicken within the avian alignment. D: Percent of various regions within the human genome mappable to each ancestral genome reconstructed along the path leading from human to the root. The positions of selected ancestors are labeled by dotted lines to indicate useful taxonomic reference points as context. E: Similar to D, but for the path of reconstructed ancestors between chicken and the root.

**Table 3:** Aggregate statistics for the 600-way alignment. The increase in computational work for the mammal alignment over the bird alignment is largely caused by the increase in the pairwise alignment phase runtime, because it scales quadratically with the size of the genomes being aligned.

| Alignment | # of genomes | Total bases | Instance-hours | Core-hours | Common ancestor size |
| --- | --- | --- | --- | --- | --- |
| 200 Mammals | 242 | 669 billion | 68,166 | 1.9 million | 1.73 Gb |
| Bird 10K | 363 | 400 billion | 5,302 | 0.2 million | 1.13 Gb |
| Combined | 605 | 1.07 trillion | 73,692 | 2.1 million | 181 Mb |

Coverage within the 600-way alignment unsurprisingly closely tracks phylogenetic distance and genome size, with e.g. a median coverage on human of 2.3 Gb from Euarchonta species, vs. 1.2 Gb from Laurasiatheria species and 1.0 Gb from Glires species (Figure 4B,C). The ancestral reconstructions within the 600-way alignment are highly complete, especially for conserved sequence: 86% of human coding bases are represented in our reconstruction of the ancestor of all placental mammals, while 95% of chicken coding bases are represented in our reconstruction of the common ancestor of avians (Figure 4D,E). The ancestral assemblies consistently contain a relatively higher proportion of avian than mammal sequence even across similar phylogenetic distance, reflecting a much more conservative mode of genome evolution in avians as well as the lower repeat content and denser gene content of avian genomes [42].

The reference-free nature of Cactus alignments enables examining genome evolution along all branches equally well, rather than being restricted to sequence present in one reference genome. In addition, the ancestral reconstructions implicitly provide a history of substitution, indel, and rearrangement events. Though this history is by its nature only a hypothetical reconstruction of the true history of genome evolution along the tree, it is by and large accurate enough to be useful. To demonstrate the utility of the indel history, we examined rates of small (⩽ 20 bp) insertion and deletion events in the 600-way alignment. As expected given previous studies [3, 16], the rate of small indels in any given branch was correlated with the rate of nucleotide substitution (an *R*^2^ of 0.49 for insertions and 0.59 for deletions), though remained much lower (1.3% of the substitution rate for insertions, and 1.7% of the substitution rate for deletions) (Figure 5A). The ancestral assemblies also represent even difficult-to-align regions such as transposable elements. We ran RepeatMasker [38] on several human ancestors, focusing on the recently-emerged L1PA6 family of L1 retrotransposons. When ascending the primate tree, approaching the origin of modern L1PA6 elements above the human-rhesus ancestor, L1PA6 elements appear increasingly similar to their consensus sequence (Figure 5B).

**Figure 5:**
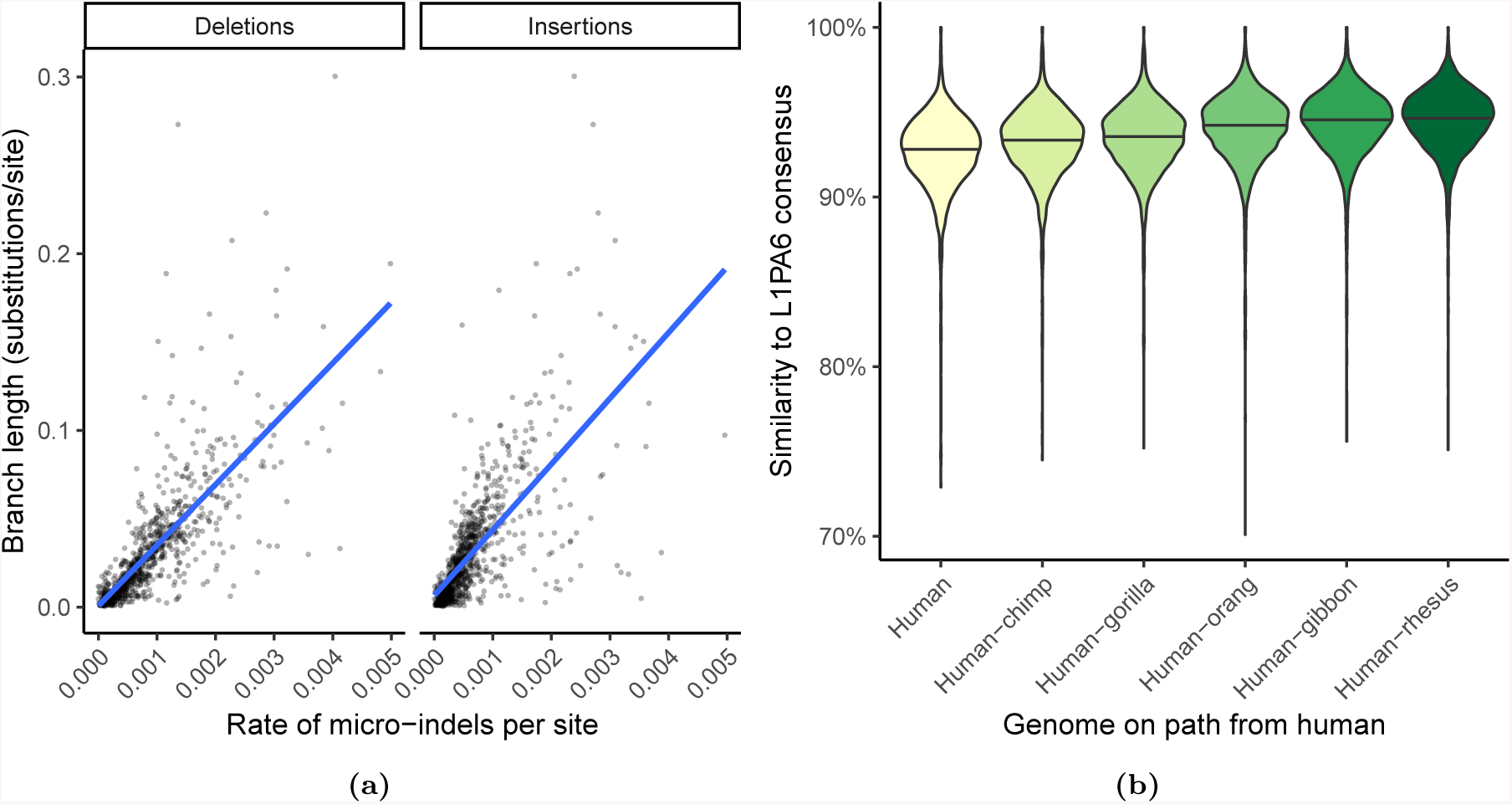
A: Rates of micro-insertions and -deletions (micro-indels) along each branch within the 600-way, compared to conventional substitutions/site branch length. B: Violin plot showing the increasing similarity to consensus of L1PA6 elements within reconstructed ancestral genomes along the path to the emergence of modern L1PA6 elements (in the human-rhesus ancestor).

The Bird 10K species were also separately aligned using MULTIZ [2] using the chicken genome as the reference, allowing us to make a comparison between the two resulting alignments. Cactus aligned more total bases to chicken than MULTIZ (an average of 69.4% of the chicken genome compared to an average of 64.9%, for an average increase of 47 Mb). Since, unlike Cactus, MULTIZ is reference-biased, the difference is more stark when looking at the number of bases aligned to a genome not used as the MULTIZ reference (an average of 79% of the zebra finch covered vs. 49.2%, for an average increase of 367Mb: see Figure 6).

**Figure 6:**
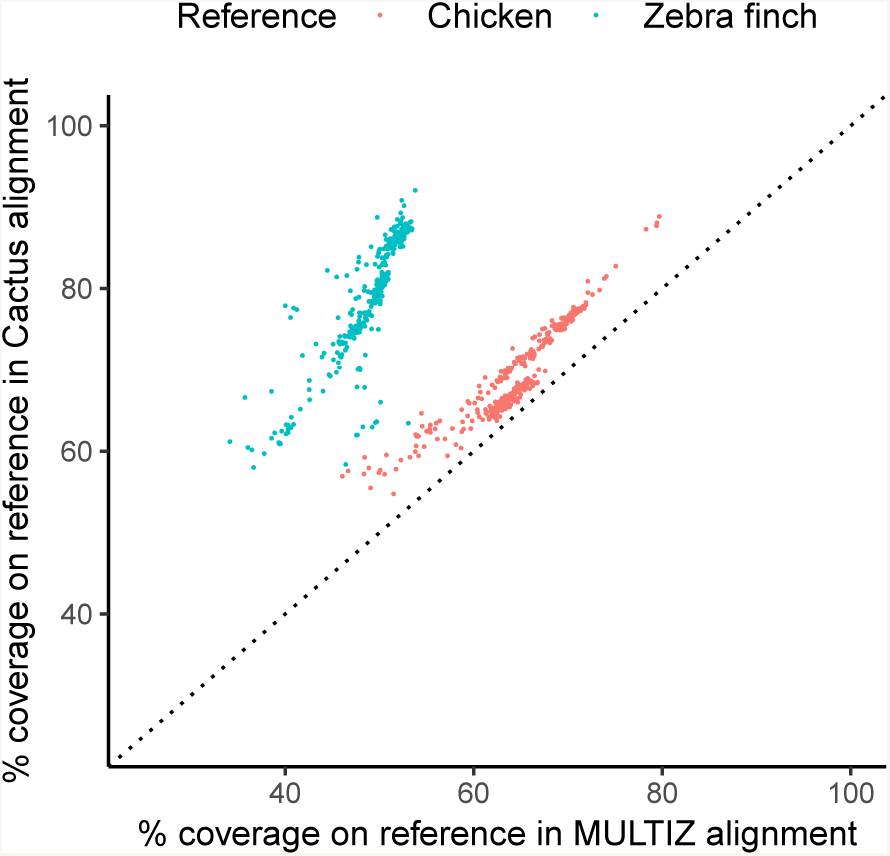
A comparison of coverage in the Cactus avian alignment compared to a chicken-referenced MULTIZ [2] alignment of the same genomes. Coverage of both alignments on chicken and zebra finch is shown to illustrate the effects of reference-bias on the completeness of the MULTIZ alignment.

## 3 Discussion

A few ambitious comparative genomics projects are already producing assemblies at the scale of tens to hundreds of species, and we anticipate that this scale of data will become much more common in the coming years. However, without a genome alignment it is impossible to relate these assemblies, and making an accurate genome alignment that large is difficult. We have demonstrated that Cactus can create alignments of hundreds of large genomes efficiently by producing an alignment relating over a trillion bases total. With this new development, we not only enable high-quality genome alignments for these projects, but also hope to set the stage for analysis of thousands to tens-of-thousands of genomes in the near future.

Furthermore, as long-read technologies become cheaper and more widely accessible, assembly quality has been rising. The age of having only a few high-quality vertebrate assemblies, like human or mouse, is at its end. As more assemblies converge on the gold-standard, “reference” level of quality displayed by GRCh38 and GRCm38, a reference-free genome alignment becomes increasingly useful. A reference-biased alignment forces the user to view genome evolution through the lens of a single, distant reference. As the average assembly becomes ever more complete and accurate, this missed opportunity to analyze regions not present in the reference grows even worse as more data is ignored and does not contribute to the alignment. For this reason, we provide a reference-free alignment, allowing analysis of genome evolution throughout the entire tree rather than in comparison to one anointed reference.

Cactus is also useful for comparison between assemblies of the same species, not just comparison between species. Often a sequencing effort will produce multiple *de novo* assemblies from different individuals, or diploid assemblies from a single individual. Alignments of these assemblies are essential for many analyses, e.g. annotation of *de novo* assemblies [11]. Cactus is easily capable of capturing even the most complex structural variation, such as copy number variation, between these assemblies.

Producing a genome alignment has usually been an arcane task, where parameters used to produce, chain, or filter the input local alignments can have an under-appreciated effect on the result. We provide Cactus as an integrated pipeline that can be used across many different compute environments, but especially thrives on modern cloud environments. It intelligently adjusts alignment parameters to maximize efficiency and accuracy depending on evolutionary distance. While genome alignment is a computationally intensive task, we have broken up the problem into small pieces that can work in heterogeneous clusters, playing to the advantages of both cheap CPU-rich machines and more expensive memory-rich machines.

We have used Cactus to produce a 600-way alignment, which is, to our knowledge, the largest-yet genome alignment of vertebrates. This alignment is already proving useful for further downstream analysis. The Bird 10K [41] and 200 Mammals [14] consortia plan to use the alignment to analyze selection at unprecedented detail across avians and mammals, respectively.

In the quest to make Cactus more efficient, optimizing the local alignment phase would offer the most return because the computational cost of the alignment is dominated by the generation of local alignments. Some less-sensitive local alignment programs are naturally more efficient than LASTZ, which is tuned for high sensitivity and long evolutionary distances. Making the local alignment phase a “pluggable” module, in which methods of generating the local alignments, or even the initial sequence graph, could be easily swapped out would be a fruitful avenue for experimentation. Cactus could potentially transition between using a less sensitive local aligner for closely related sequence and a more sensitive aligner across long evolutionary distance, much in the same way that we change alignment parameters based on evolutionary distance today.

As alignments become larger and more expensive to compute, it becomes much more important to be able to update them (by e.g. adding a new genome or updating an assembly) without recomputing the entire alignment. Cactus’s progressive alignment framework, combined with special functionality in the HAL toolkit [18] makes it possible to make these changes very efficiently: costing only a single subproblem’s worth of computation time, usually about 120 CPU days. However, there is currently an appreciable amount of manual work involved in the process of adding, removing, or updating an assembly within an existing alignment. Making this simpler and more automated would be an interesting future direction, one that would potentially allow a very large alignment resource to be used and updated for years, with collaborators adding in their genomes of interest cost-effectively.

## 4 Data availability

The 600-way alignment is available in HAL format at https://alignment-output.s3.amazonaws.com/600way.hal. We anticipate that because of the large file size and the relatively small amount of alignable sequence between avians and mammals, most users will be interested in the alignments of the avian and mammalian clades. For this reason, we also provide the subset of the alignment containing the 200 Mammals genomes at https://alignment-output.s3.amazonaws.com/200m-v1.hal, and the subset of the alignment containing the Bird 10K genomes at https://alignment-output.s3.amazonaws.com/birds-final.hal.

A visualization of the alignments and associated data is available by loading our assembly hub into the UCSC browser. By copying the hub link https://comparative-genomics-hubs.s3-us-west-2.amazonaws.com/600way_hub.txt into the “Track Hubs” page, the 605 genomes and associated tracks will be available.

## 5 Code availability

The Cactus pipeline is available at https://github.com/ComparativeGenomicsToolkit/cactus. The exact version of Cactus used for each of the analyses described above varies; the commit used in each analysis are available in the supplementary material.

## Supporting information

Supplementary Material: Section S1, Figures S1-5, Tables S1-3

Guide trees for 48-bird guide-tree analysis

## 6 Acknowledgements

Research reported in this publication was supported by the National Human Genome Research Institute of the National Institutes of Health under Award Numbers 1R01HG008742, R01HG009737, U54HG007990 and 2U41HG007234. Research reported in this publication was supported by the National Heart, Lung, And Blood Institute of the National Institutes of Health under Award Number U01HL137183. The content is solely the responsibility of the authors and does not necessarily represent the official views of the National Institutes of Health. 7 Methods 7.1

## 7 Methods

### 7.1 Cactus

#### 7.1.1 Preliminary repeat-masking

Cactus requires input genomes to be soft-masked, but often repetitive sequence goes unmasked due to poor masking or incomplete repeat libraries for newly-sequenced species. This can negatively affect alignment runtimes (as alignments need to be enumerated to and from all copies of a repetitive sequence) and impact quality. For this reason, we mask overabundant sequence before alignment, using a strategy not based on alignment to repeat consensus libraries, but on over-representation of alignments. We first divide each genome into small, mutually overlapping chunks. For each chunk, we align it to itself and a configurable amount of other randomly sampled chunks (currently 20% of the total pool). To avoid combinatorial explosion due to unmasked repetitive sequence, we use a special mode of LASTZ [17] which stops exploring alignments from any region early if a maximum depth is reached. We then soft-mask any region covered by more than a certain configurable number of these alignments (currently set to 50).

#### 7.1.2 Local alignment and outgroup selection

The alignment process for each subproblem begins with a series of local alignments generated using LASTZ [17]. The local alignments fall into two sets: a set of all-against-all alignments among the ingroup genomes, and a set of alignments from ingroup genomes to outgroup genomes.

We have found outgroup selection to be absolutely crucial in creating an acceptable ancestral reconstruction: any missing data or misassembly in the outgroup that causes a true deletion in one of the ingroups to be misinterpreted as an insertion in others will mean that the ancestor contains less sequence than it ought to. This missing sequence in turn impacts the alignment between the entire subtree below the reconstructed ancestor and the entire supertree above it: the missing sequence will never be aligned between the subtree and supertree. To avoid this we attempt to use multiple outgroup genomes in each subproblem (3 by default). Naively aligning each ingroup against multiple outgroups would significantly increase the computation time; to avoid this we note that in general any region already containing an outgroup alignment benefits very little from aligning an additional outgroup. Therefore, we iteratively align each ingroup against one outgroup at a time, pruning away any ingroup sequence already covered by the previous outgroup alignments. In this way the computational cost is reduced to be far less than naively aligning against the entire outgroup set, while still retaining nearly all of the benefit. In addition, we allow the user to designate certain genomes in the input as being of particularly high quality; these are chosen as outgroups if possible to avoid problems with missing data in regions like mitochondrial or sex chromosomes that are often missing from some assemblies but not others.

#### 7.1.3 Ancestral genome reconstruction

The core of what makes the progressive alignment algorithm possible is the ancestral reconstruction generated in each subproblem. This assembly serves as a summary of each subproblem alignment; the alignable sequence between the genomes in the subtree below the ancestor, as well as that alignable between the subtree and the supertree above the ancestor, is all present in the ancestral reconstruction. The ancestral sequence contains a base for each column in all blocks which contain an alignment between two ingroups and/or an ingroup and an outgroup — any alignment purely between outgroups is discarded. The order and orientation of the blocks relative to one another is chosen via a previously published algorithm for ordering a pangenome [34].

The identity of the ancestral bases is inferred via maximum-likelihood on a Jukes-Cantor model [24] of evolution using Felsenstein’s pruning algorithm [9] on the subtree of the guide tree induced by the genomes in the subproblem. These base-calls are generated as the alignment is being made, so they necessarily take only a part of the alignment information into account and may be different than the ideal base-calls would be if taking into account information across the entire alignment. However, we provide a tool, `ancestorsML`, distributed as part of the HAL toolkit [18], that re-estimates ancestral base-calls after completion of the alignment if desired.

#### 7.1.4 Paralogy resolution

Previous beta versions of progressive Cactus relied on an outgroup-based heuristic to resolve duplication timing. This heuristic, which we term “single-copy outgroup filtering”, separated collections of ingroup regions based on their similarity to outgroup regions, ensuring that at most one outgroup region could be present per block: the one most similar to the block’s ingroup sequences. Assuming that the outgroup contains the proper number of copies and each ingroup copy is indeed most similar to an orthologous outgroup copy, this should function correctly. However, this method is very sensitive to incomplete outgroup assemblies (containing an incorrect number of copies of a duplicated region) or variation in similarity between closely related paralogs causing assignment to the wrong copy. As seen in Figure 3, this filtering method tended to resolve duplications far too early, often causing paralogs to be called as orthologs (for example, implicitly labeling 6.1% of human sequence as duplicated along the chimpanzee lineage, which is certainly an overestimate).

To remedy this problem, we developed an improved duplication-timing method, which we termed “best-hit filtering” in the earlier text. The method assigns, for every base in every input genome, a single *primary* pairwise alignment (the highest-scoring alignment involving that base, if it has been aligned) and a set of *secondary* pairwise alignments (all others involving that base). All primary alignments are added to the initial graph unconditionally, as they represent the most likely ortholog relationship (or in the case of multiple orthology, likely a random ortholog) (Figure S5). The set of primary alignments represents a conservative set of alignment relationships that should include nearly no alignments to ancient paralogs. However, in regions that have undergone many rounds of lineage-specific duplications (which should all be aligned together in the restricted duplication-timing problem we describe above), the set of primary alignments will often by chance not align all copies together. For this reason, we also allow some of the secondary alignments into the initial graph, after adding the primaries, though with additional restrictions because the secondary alignments will inevitably contain some alignments to distant paralogs. We only allow in those secondary alignments that would not merge two existing blocks that both contain sequences from multiple species — this allows lineage-specific duplications to correctly land in the same block, while avoiding merging blocks from likely-paralogous alignments.

#### 7.1.5 Removing recoverable chains

Due to the insensitive and approximate nature of the input local alignments, homologies are often missed in the input to the CAF algorithm. Alignment blocks that are “incomplete”, i.e. contain some true homologies but miss others, can cause issues for the CAF algorithm: if these incomplete blocks make it into the output Cactus graph, the missing homologies can never be recovered by the BAR algorithm because to preserve the structure of the cactus graph, BAR cannot modify existing alignment blocks, only add new ones. To remedy this issue, we developed a method to remove likely-incomplete blocks as part of the algorithm, which we term “removing recoverable chains”. In short, this method runs as a post-processing step to the original CAF algorithm, removing blocks which contain only homologies that could recovered by the BAR algorithm extending from neighboring blocks. Adding this post-filtering step noticeably increases coverage, especially for distant genomes in large trees (Figure S4). For further detail on the process, see Section S1.8.

### 7.2 Adding a new genome to an existing alignment

There are three possible ways to add a genome into an existing alignment, depending on the desired phylogenetic position of the genome. Adding a genome as an outgroup is straightforward, since it follows the normal progressive process: the root of the existing alignment and the new genome can be aligned together into a supertree alignment, which the existing subtree alignment can be appended to. A genome can be added as a new child of an existing internal node by simply aligning the new child, its siblings, and its parent together, without inferring a new ancestral genome. Adding a genome by splitting an existing branch is the least straightforward, but is key if the topology of the alignment or the accuracy of the ancestral genomes is important. To add a genome to an existing alignment, two subproblems are required: one relating the new genome and its new sibling in the target tree, constructing the ancestral genome that will split the existing branch, and one relating this new ancestral genome, its sibling, and its parent.

After addition of a new genome as an ingroup (by adding it to a node or a branch), at most a single ancestral sequence is re-inferred. This prevents any information from the new genome from propagating to the rest of the tree. While this saves significant effort in recomputing other parts of the alignment, it also means that, occasionally, rare stretches of sequence in a newly added genome would not be properly aligned to distant outgroups because they were deleted or missing in the new genome’s close relatives. Re-inferring the ancestral genomes on the path from newly added genomes to the root should address this issue if it appears.

## References

[1] J. Armstrong, I. T. Fiddes, M. Diekhans, and B. Paten. Whole-Genome Alignment and Comparative Annotation. Annu Rev Anim Biosci, Oct 2018.

[2] Mathieu Blanchette, W. James Kent, Cathy Riemer, Laura Elnitski, Arian F A Smith, Krishna M. Roskin, Robert Baertsch, Kate Rosenbloom, Hiram Clawson, Eric D. Green, David Haussler, and Webb Miller. Aligning multiple genomic sequences with the threaded blockset aligner. Genome Research, 14(4):708–715, 2004.

[3] Jian-Qun Chen, Ying Wu, Haiwang Yang, Joy Bergelson, Martin Kreitman, and Dacheng Tian. Variation in the Ratio of Nucleotide Substitution and Indel Rates across Genomes in Mammals and Bacteria. Molecular Biology and Evolution, 26(7):1523–1531, 03 2009.

[4] Aaron E. Darling, Bob Mau, and Nicole T. Perna. Progressivemauve: Multiple genome alignment with gene gain, loss and rearrangement. PLoS ONE, 5(6), 2010.

[5] P. Dobrynin, S. Liu, G. Tamazian, Z. Xiong, A. A. Yurchenko, K. Krasheninnikova, S. Kliver, A. Schmidt-Kuntzel, K. P. Koepfli, W. Johnson, L. F. Kuderna, R. Garcia-Perez, M. d. Manuel, R. Godinez, A. Komissarov, A. Makunin, V. Brukhin, W. Qiu, L. Zhou, F. Li, J. Yi, C. Driscoll, A. Antunes, T. K. Oleksyk, E. Eizirik, P. Perelman, M. Roelke, D. Wildt, M. Diekhans, T. Marques-Bonet, L. Marker, J. Bhak, J. Wang, G. Zhang, and S. J. O’Brien. Genomic legacy of the African cheetah, Acinonyx jubatus. Genome Biol., 16:277, Dec 2015.

[6] Dent Earl, Ngan Nguyen, Glenn Hickey, Robert S Harris, Stephen Fitzgerald, Kathryn Beal, Igor Seledtsov, Vladimir Molodtsov, Brian J Raney, Hiram Clawson, et al. Alignathon: a competitive assessment of whole-genome alignment methods. Genome research, 24(12):2077–2089, 2014.

[7] R. C. Edgar, G. Asimenos, S. Batzoglou, and A. Sidow. Evolver: a whole-genome sequence evolution simulator. https://www.drive5.com/evolver.

[8] J. Eid, A. Fehr, J. Gray, K. Luong, J. Lyle, G. Otto, P. Peluso, D. Rank, P. Baybayan, B. Bettman, A. Bibillo, K. Bjornson, B. Chaudhuri, F. Christians, R. Cicero, S. Clark, R. Dalal, A. Dewinter, J. Dixon, M. Foquet, A. Gaertner, P. Hardenbol, C. Heiner, K. Hester, D. Holden, G. Kearns, X. Kong, R. Kuse, Y. Lacroix, S. Lin, P. Lundquist, C. Ma, P. Marks, M. Maxham, D. Murphy, I. Park, T. Pham, M. Phillips, J. Roy, R. Sebra, G. Shen, J. Sorenson, A. Tomaney, K. Travers, M. Trulson, J. Vieceli, J. Wegener, D. Wu, A. Yang, D. Zaccarin, P. Zhao, F. Zhong, J. Korlach, and S. Turner. Real-time DNA sequencing from single polymerase molecules. Science, 323(5910):133–138, Jan 2009.

[9] Joseph Felsenstein. Maximum Likelihood and Minimum-Steps Methods for Estimating Evolutionary Trees from Data on Discrete Characters. Systematic Biology, 22(3):240–249, 09 1973.

[10] D F Feng and R F Doolittle. Progressive sequence alignment as a prerequisite to correct phylogenetic trees. Journal of molecular evolution, 25(4):351–360, 1987.

[11] Ian T Fiddes, Joel Armstrong, Mark Diekhans, Stefanie Nachtweide, Zev N Kronenberg, Jason G Underwood, David Gordon, Dent Earl, Thomas Keane, Evan E Eichler, et al. Comparative annotation toolkit (cat)-simultaneous clade and personal genome annotation. bioRxiv, page 231118, 2017.

[12] Manuel Garber, Mitchell Guttman, Michele Clamp, Michael C. Zody, Nir Friedman, and Xiaohui Xie. Identifying novel constrained elements by exploiting biased substitution patterns. Bioinformatics, 25(12):54–62, 2009.

[13] E. Garrison, J. Siren, A. M. Novak, G. Hickey, J. M. Eizenga, E. T. Dawson, W. Jones, S. Garg, C. Markello, M. F. Lin, B. Paten, and R. Durbin. Variation graph toolkit improves read mapping by representing genetic variation in the reference. Nat. Biotechnol., 36(9):875–879, 10 2018.

[14] D. Genereux, J. Johnson, V. Marinescu, E. Murén, J. Armstrong, A. S. Armero, D. Juan, G. Bejerano, N. Casewell, L. Chemnick, J. Damas, F. de Palma, M. Diekhans, I. Fiddes, M. Garber, L. Goodman, W. Haerty, M. Houck, R. Hubley, T. Kivioja, L. Kuderna, E. Lander, Marques-Bonet T., J. Meadows, W. Murphy, W. Nash, H. J. Noh, M. Nweeia, B. Paten, A. Pfenning, K. Pollard, D. Ray, B. Shapiro, A. Smit, M. Springer, C. Steiner, R. Swofford, J. Taipale, E. Teeling, J. Turner-Maier, K. Lewin, J. Alfoldi, O. Ryder, B. Birren, and K. Lindblad-Toh. Genomics in an age of extinction. in submission.

[15] David Gordon, John Huddleston, Mark JP Chaisson, Christopher M Hill, Zev N Kronenberg, Katherine M Munson, Maika Malig, Archana Raja, Ian Fiddes, LaDeana W Hillier, et al. Long-read sequence assembly of the gorilla genome. Science, 352(6281):aae0344, 2016.

[16] R. E. Green, E. L. Braun, J. Armstrong, D. Earl, N. Nguyen, G. Hickey, M. W. Vandewege, J. A. St. John, S. Capella-Gutierrez, T. A. Castoe, C. Kern, M. K. Fujita, J. C. Opazo, J. Jurka, K. K. Kojima, J. Caballero, R. M. Hubley, A. F. Smit, R. N. Platt, C. A. Lavoie, M. P. Ramakodi, J. W. Finger, A. Suh, S. R. Isberg, L. Miles, A. Y. Chong, W. Jaratlerdsiri, J. Gongora, C. Moran, A. Iriarte, J. McCormack, S. C. Burgess, S. V. Edwards, E. Lyons, C. Williams, M. Breen, J. T. Howard, C. R. Gresham, D. G. Peterson, J. Schmitz, D. D. Pollock, D. Haussler, E. W. Triplett, G. Zhang, N. Irie, E. D. Jarvis, C. A. Brochu, C. J. Schmidt, F. M. McCarthy, B. C. Faircloth, F. G. Hoffmann, T. C. Glenn, T. Gabaldon, B. Paten, and D. A. Ray. Three crocodilian genomes reveal ancestral patterns of evolution among archosaurs. Science, 346(6215):1254449–1254449, 2014.

[17] R.S. Harris. Improved pairwise alignment of genomic DNA. PhD thesis, The Pennsylvania State University, 2007.

[18] Glenn Hickey, Benedict Paten, Dent Earl, Daniel Zerbino, and David Haussler. Hal: a hierarchical format for storing and analyzing multiple genome alignments. Bioinformatics, page btt128, 2013.

[19] Melissa J. Hubisz, Katherine S. Pollard, and Adam Siepel. PHAST and RPHAST: Phylogenetic analysis with space/time models. Briefings in Bioinformatics, 12(1):41–51, 2011.

[20] S. Huntley, D. M. Baggott, A. T. Hamilton, M. Tran-Gyamfi, S. Yang, J. Kim, L. Gordon, E. Branscomb, and L. Stubbs. A comprehensive catalog of human KRAB-associated zinc finger genes: insights into the evolutionary history of a large family of transcriptional repressors. Genome Res., 16(5):669–677, May 2006.

[21] M. Jain, S. Koren, K. H. Miga, J. Quick, A. C. Rand, T. A. Sasani, J. R. Tyson, A. D. Beggs, A. T. Dilthey, I. T. Fiddes, S. Malla, H. Marriott, T. Nieto, J. O’Grady, H. E. Olsen, B. S. Pedersen, A. Rhie, H. Richardson, A. R. Quinlan, T. P. Snutch, L. Tee, B. Paten, A. M. Phillippy, J. T. Simpson, N. J. Loman, and M. Loose. Nanopore sequencing and assembly of a human genome with ultra-long reads. Nat. Biotechnol., 36(4):338–345, 04 2018.

[22] M. Jain, H. E. Olsen, B. Paten, and M. Akeson. The Oxford Nanopore MinION: delivery of nanopore sequencing to the genomics community. Genome Biol., 17(1):239, 11 2016.

[23] Erich D Jarvis, Siavash Mirarab, Andre J Aberer, Bo Li, Peter Houde, Cai Li, Simon YW Ho, Brant C Faircloth, Benoit Nabholz, Jason T Howard, et al. Whole-genome analyses resolve early branches in the tree of life of modern birds. Science, 346(6215):1320–1331, 2014.

[24] Thomas H Jukes and Charles R Cantor. Evolution of protein molecules. Mammalian protein metabolism, 3(21):132, 1969.

[25] P. A. Kitts, D. M. Church, F. Thibaud-Nissen, J. Choi, V. Hem, V. Sapojnikov, R. G. Smith, aT. Tatusova, C. Xiang, A. Zherikov, M. DiCuccio, T. D. Murphy, K. D. Pruitt, and A. Kimchi. Assembly: a resource for assembled genomes at NCBI. Nucleic Acids Res., 44(D1):73–80, Jan 2016.

[26] Klaus-Peter Koepfli, Benedict Paten, the Genome 10K Community of Scientists, and Stephen J. O’Brien. The genome 10k project: A way forward. Annual Review of Animal Biosciences, 3(1):57–111, 2015. PMID: 25689317.

[27] Jonas Korlach, Gregory Gedman, Sarah B. Kingan, Chen-Shan Chin, Jason T. Howard, Jean-Nicolas Audet, Lindsey Cantin, and Erich D. Jarvis. De novo PacBio long-read and phased avian genome assemblies correct and add to reference genes generated with intermediate and short reads. GigaScience, 6(10), 08 2017.

[28] Z. N. Kronenberg, I. T. Fiddes, D. Gordon, S. Murali, S. Cantsilieris, O. S. Meyerson, J. G. Underwood, B. J. Nelson, M. J. P. Chaisson, M. L. Dougherty, K. M. Munson, A. R. Hastie, M. Diekhans, F. Hormozdiari, N. Lorusso, K. Hoekzema, R. Qiu, K. Clark, A. Raja, A. E. Welch, M. Sorensen, C. Baker, R. S. Fulton, J. Armstrong, T. A. Graves-Lindsay, A. M. Denli, E. R. Hoppe, P. Hsieh, C. M. Hill, A. W. C. Pang, J. Lee, E. T. Lam, S. K. Dutcher, F. H. Gage, W. C. Warren, J. Shendure, D. Haussler, V. A. Schneider, H. Cao, M. Ventura, R. K. Wilson, B. Paten, A. Pollen, and E. E. Eichler. High-resolution comparative analysis of great ape genomes. Science, 360(6393), 06 2018.

[29] Gregory M. Kurtzer, Vanessa Sochat, and Michael W. Bauer. Singularity: Scientific containers for mobility of compute. PLoS ONE, 12(5):1–20, 2017.

[30] S. König, L. W. Romoth, L. Gerischer, and M. Stanke. Simultaneous gene finding in multiple genomes. Bioinformatics, 32(22):3388–3395, 11 2016.

[31] Harris A. Lewin, Gene E. Robinson, W. John Kress, William J. Baker, Jonathan Coddington, Keith A. Crandall, Richard Durbin, Scott V. Edwards, Félix Forest, M. Thomas P. Gilbert, Melissa M. Goldstein, Igor V. Grigoriev, Kevin J. Hackett, David Haussler, Erich D. Jarvis, Warren E. Johnson, Aristides Patrinos, Stephen Richards, Juan Carlos Castilla-Rubio, Marie-Anne van Sluys, Pamela S. Soltis, Xun Xu, Huanming Yang, and Guojie Zhang. Earth biogenome project: Sequencing life for the future of life. Proceedings of the National Academy of Sciences, 115(17):4325–4333, 2018.

[32] J. Lilue, A. G. Doran, I. T. Fiddes, M. Abrudan, J. Armstrong, R. Bennett, W. Chow, J. Collins, S. Collins, A. Czechanski, P. Danecek, M. Diekhans, D. D. Dolle, M. Dunn, R. Durbin, D. Earl, A. Ferguson-Smith, P. Flicek, J. Flint, A. Frankish, B. Fu, M. Gerstein, J. Gilbert, L. Goodstadt, J. Harrow, K. Howe, X. Ibarra-Soria, M. Kolmogorov, C. J. Lelliott, D. W. Logan, J. Loveland, C. E. Mathews, R. Mott, P. Muir, S. Nachtweide, F. C. P. Navarro, D. T. Odom, N. Park, S. Pelan, S. K. Pham, M. Quail, L. Reinholdt, L. Romoth, L. Shirley, C. Sisu, M. Sjoberg-Herrera, M. Stanke, C. Steward, M. Thomas, G. Threadgold, D. Thybert, J. Torrance, K. Wong, J. Wood, B. Yalcin, F. Yang, D. J. Adams, B. Paten, and T. M. Keane. Sixteen diverse laboratory mouse reference genomes define strain-specific haplotypes and novel functional loci. Nat. Genet., 50(11):1574–1583, Nov 2018.

[33] Ngan Nguyen, Glenn Hickey, Brian J. Raney, Joel Armstrong, Hiram Clawson, Ann Zweig, Donna Karolchik, William James Kent, David Haussler, and Benedict Paten. Comparative assembly hubs: Web-accessible browsers for comparative genomics. Bioinformatics, 30(23):3293–3301, 2014.

[34] Ngan Nguyen, Glenn Hickey, Daniel R Zerbino, Brian Raney, Dent Earl, Joel Armstrong, W James Kent, David Haussler, and Benedict Paten. Building a pan-genome reference for a population. Journal of computational biology: a journal of computational molecular cell biology, 22(5):387–401, 2015.

[35] Benedict Paten, Dent Earl, Ngan Nguyen, Mark Diekhans, Daniel Zerbino, and David Haussler. Cactus: Algorithms for genome multiple sequence alignment. Genome research, 21(9):1512–1528, 2011.

[36] M. N. Price, P. S. Dehal, and A. P. Arkin. FastTree 2–approximately maximum-likelihood trees for large alignments. PLoS ONE, 5(3):e9490, Mar 2010.

[37] R. O. Prum, J. S. Berv, A. Dornburg, D. J. Field, J. P. Townsend, E. M. Lemmon, and A. R. Lemmon. A comprehensive phylogeny of birds (Aves) using targeted next-generation DNA sequencing. Nature, 526(7574):569–573, Oct 2015.

[38] Smit, A. F. A. and Hubley, R. and Green, P. RepeatMasker Open-4.0. http://www.repeatmasker.org, 2013-2015.

[39] John Vivian, Arjun Arkal Rao, Frank Austin Nothaft, Christopher Ketchum, Joel Armstrong, Adam Novak, Jacob Pfeil, Jake Narkizian, Alden D Deran, Audrey Musselman-Brown, et al. Toil enables reproducible, open source, big biomedical data analyses. Nature biotechnology, 35(4):314, 2017.

[40] N. I. Weisenfeld, V. Kumar, P. Shah, D. M. Church, and D. B. Jaffe. Direct determination of diploid genome sequences. Genome Res., 27(5):757–767, 05 2017.

[41] G. Zhang, C. Rahbek, G. R. Graves, F. Lei, E. D. Jarvis, and M. T. Gilbert. Genomics: Bird sequencing project takes off. Nature, 522(7554):34, Jun 2015.

[42] Guojie Zhang, Cai Li, Qiye Li, Bo Li, Denis M Larkin, Chul Lee, Jay F Storz, Agostinho Antunes, Matthew J Greenwold, Robert W Meredith, et al. Comparative genomics reveals insights into avian genome evolution and adaptation. Science, 346(6215):1311–1320, 2014.

