## Supplementary Material: Section S1, Figures S1-5, Tables S1-3 for "Progressive alignment with Cactus: a multiple-genome aligner for the thousand-genome era"

### S1 Supplementary Methods

#### S1.1 Evaluation on simulated data

20 primate genomes were simulated using Evolver [2], managed using the evolverSimControl (https://github.com/dentearl/evolverSimControl, commit b3236deb) pipeline. The root genome used was derived from 30 megabases selected from the hg19 genome, and is available at http://courtyard.gi.ucsc.edu/~jcarmstr/datastore/progressiveCactusEvolverSim.tar.gz along with the Evolver configuration files that were used. The species tree used for the simulation was obtained from a catarrhine subtree of the 100-way alignment tree available on the UCSC browser. The tree used was, in Newick format:

(((((((Human:0.00655,Chimp:0.00684)anc0e:0.00122,Bonobo:0.00784)anc1e:0.003,Gorilla:0.008964)anc2e:0.009693,Orangutan:0.01894)anc3e:0.003471,Gibbon:0.02227)anc4e:0.01204,(((((Rhesus:0.004991,Crab_eating_macaque:0.005991)anc5e:0.001,Sooty_mangabey:0.001)anc6e:0.005,Baboon:0.003042)anc7e:0.01061,(Green_monkey:0.027,Drill:0.03)anc8e:0.002)anc9e:0.003,((Proboscis_monkey:0.0007,Angolan_colobus:0.0008)anc10e:0.005,(Golden_snub-nosed_monkey:0.0007,Black_snub_nosed_monkey:0.0008)anc11e:0.004)anc12e:0.009)anc13e:0.02)anc14e:0.02183,(((Marmoset:0.03,Squirrel_monkey:0.01035)anc15e:0.01065,White-faced_sapajou:0.009)anc16e:0.01,Nancy_Mas_night_monkey:0.01)anc17e:0.01)anc18e;

The alignments were generated using Cactus commit 51eb980b. The input files (the simulated genomes as well as input files and Cactus configuration file) are available at http://courtyard.gi.ucsc.edu/~jcarmstr/datastore/progressiveCactus.EvolverSim.CactusInput.tar.gz. A non-default configuration (included in the dataset) was used to change the alignment filtering in both runs to better support the high degree of polytomy in the star-tree runs. Four sets of 2, 6, 10, and 20 genomes were used, each of which were run three times to generate runtime estimates.

The runtime statistics were gathered using the toil stats command (the overall Clock time was used, which represents CPU time spent across all jobs). To generate the recall and precision statistics, MAFs were exported for each run (using hal2maf with the --onlyOrthologs option using the rhesus genome as a reference) and compared to the Evolver MAF using mafComparator (https://github.com/dentearl/mafTools, commit 82077ac3).

#### S1.2 Adding a new genome to the simulated alignment

We evaluated the accuracy of adding a genome to an existing alignment by creating a new alignment of 19 of the 20 simulated genomes described above (holding out the "Crab_eating_macacque" genome), then adding it back in after the fact. All alignments for this analysis were generated using Cactus commit 49e80082.

To add the crab-eating macaque back in as the child of an existing node (the add-to-node strategy), we ran a single new alignment with the tree (Rhesus:0.006, Crab_eating_macaque:0.007, Sooty_mangabey:0.001)anc6e;. The anc6e genome from the original, held-out alignment was used as a unreconstructed ancestral input sequence. We set the "runMapQFiltering" option in the config file to "0" and the "alignmentFilter" option to "singleCopyOutgroup", since these options produce a better alignment of polytomies. We merged the resulting HAL file into a new copy of the existing alignment via the command halReplaceGenome<copyofheld-outalignment>anc6e--topAlignmentFile<held-outalignment>--bottomAlignmentFile<add-to-nodealignment>.

To add the macaque by splitting a branch (the add-to-branch strategy), we ran two separate alignments. We ran the first with the tree (((Rhesus:0.004991, Crab_eating_macaque:0.005991)anc5e:0.001, Sooty_mangabey:0.001)anc6e:0.005, Baboon:0.003042)anc7e; (with the --root anc5e option so that only a single subproblem was run), generating a newly reconstruted anc5e ancestor. We then ran a second alignment with the tree (anc5e:0.001, Sooty_mangabey:0.001)anc6e;, again providing the anc6e assembly from the original alignment rather than inferring a new reconstruction. (We note that these two subproblems could have been run in a single alignment invocation,

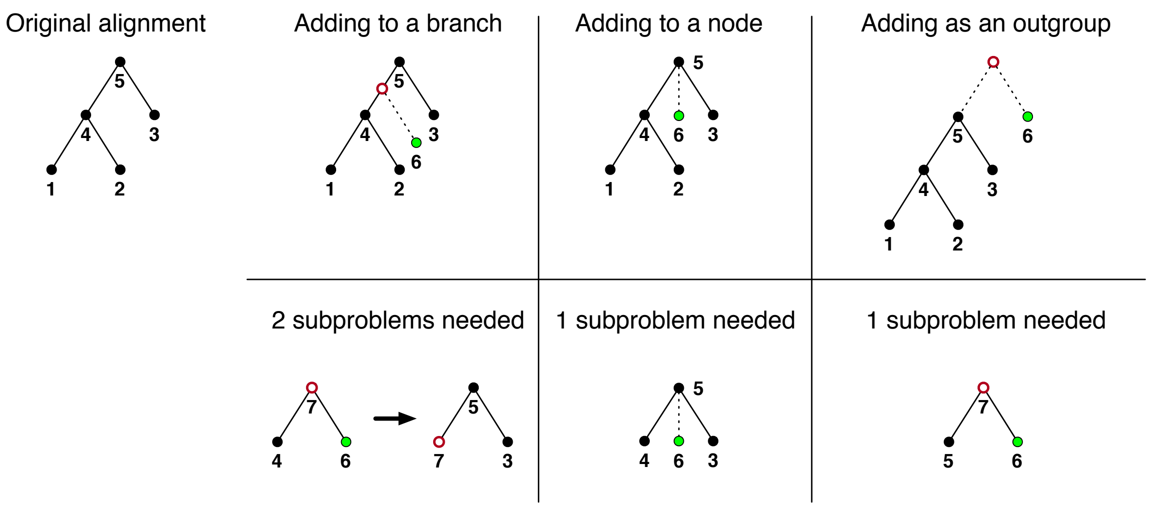


**Figure S1:** Methods of adding a genome to a Cactus alignment. The top row shows the different ways of adding a new genome given its phylogenetic position, and the bottom row shows what subproblems would need to be computed for the new genome to be properly merged into the existing alignment. Green circles represent a new genome, and red circles represent newly reconstructed genomes.

resulting in the same amount of alignment work but a slightly more complicated merging process.) To merge these two add-to-branch intermediate alignments into a full alignment, we first removed the Rhesus genome from a new copy of the held-out alignment. We then ran halAddToBranch <held-out alignment> <first add-to-branch alignment> <second add-to-branch alignment> anc6e anc5e Rhesus Crab_eating_macaque 0.001 0.006.

We evaluated the performance of these new alignments using mafComparator in the same way as described in Section S1.1. In the interest of narrowly determining accuracy of alignments involving the newly added genome, we counted only aligned pairs involving the Crab_eating_macaque genome when calculating precision, recall, and F1 scores.

#### S1.3 Evaluation of the effect of the guide tree

The guide-tree analysis was performed on a set of 48 bird genomes originally published in 2014 [7]. To reduce the amount of alignment work required, we subsetted these genomes down to the size of only a single chromosome, chicken chromosome 1 (by removing any contig or scaffold which had less than 20% of its sequence alignable to chicken chromosome 1). We used Cactus commit 36304707 for all alignments in this analysis.

The Prum and Jarvis topologies were adapted from [14] and [7], respectively. The "permuted" topology was generated starting from the Jarvis topology, via 3 randomly chosen subtree-prune-regraft operations followed by 3 random nearest-neighbor-interchange operations. Each of these three topologies had branch-length estimates performed using phyloFit from the PHAST package [6] based on fourfold-degenerate sites of BUSCO orthologs. Finally, the "Consensus" tree was produced as a strict consensus of the Jarvis and Prum trees (collapsing all groupings that were not the same in both trees) using the ape::consensus method from the APE R package [10]. The branch-lengths for this tree were generated from the fitted branch lengths for the two input trees, using the consensus.edges function of the phytools R package [15]. The four final trees that were used in the four Cactus alignments are shown in Figure S2, and available in supplementary data in Newick format.

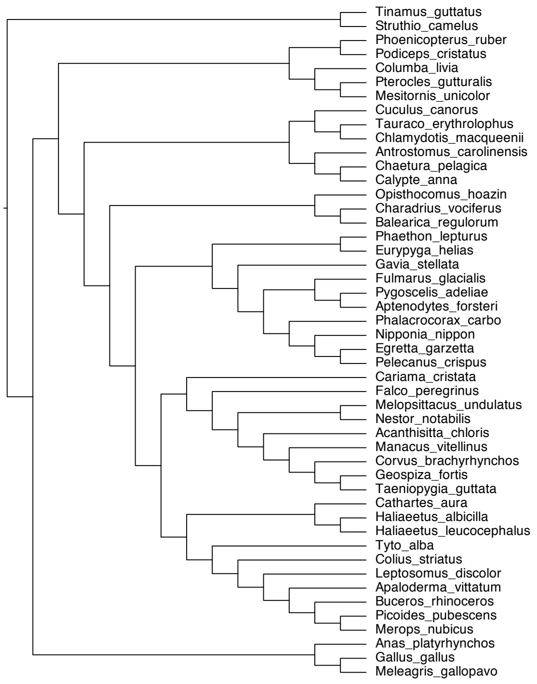


**(a)** Jarvis

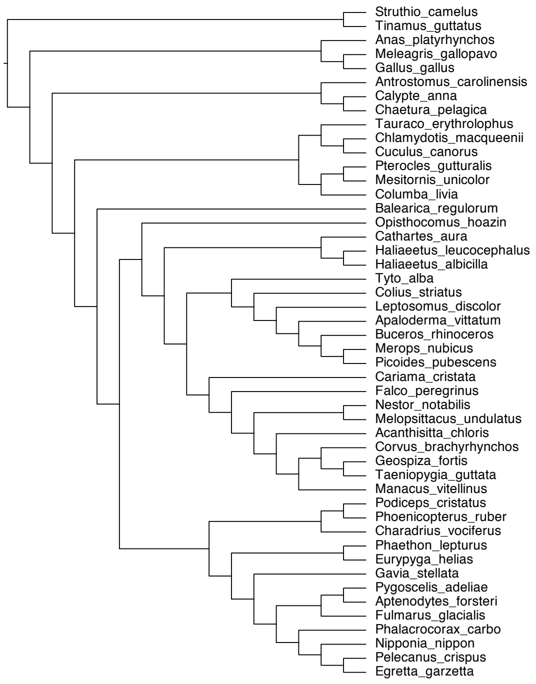


**(b)** Prum

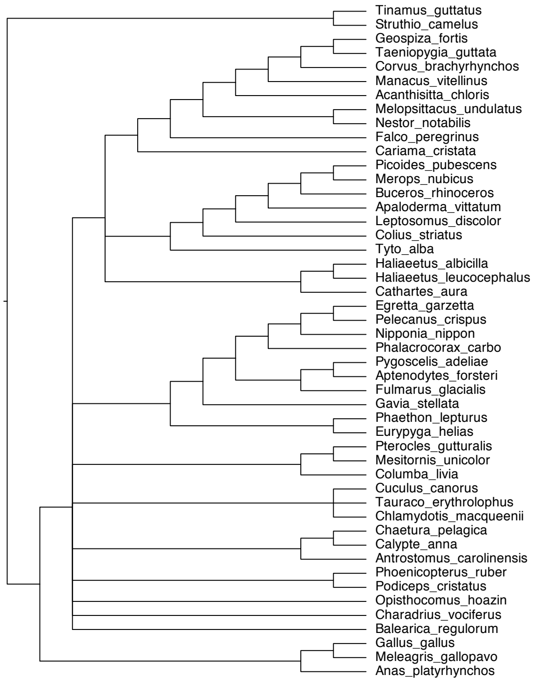


**(c)** Consensus

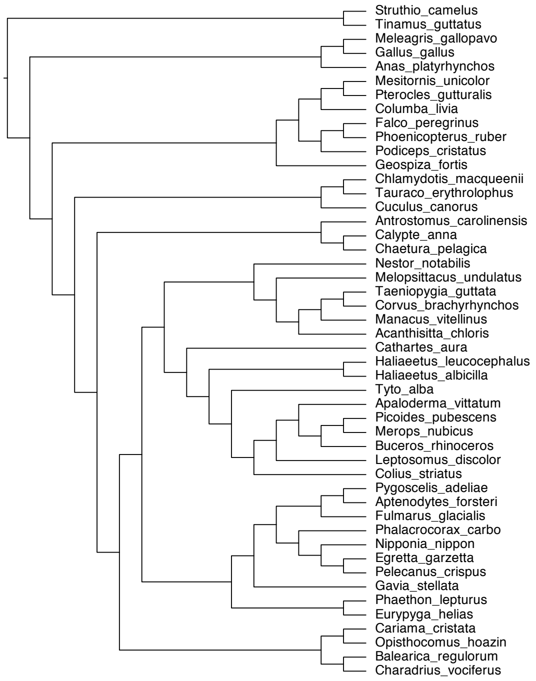


**(d)** Permuted

**Figure S2:** Guide trees used in the guide-tree influence analysis.

| Alignment | URL |
|---|---|
| Jarvis | `https://s3.amazonaws.com/alignment-output/cactus48BIRDS_jarvis14.hal` |
| Prum | `https://s3.amazonaws.com/alignment-output/cactus48BIRDS_prum15.hal` |
| Consensus | `https://s3.amazonaws.com/alignment-output/cactus48BIRDS_consensus.hal` |
| Permuted | `https://s3.amazonaws.com/alignment-output/cactus48BIRDS_permute.hal` |

**Table S1:** Alignments used in the guide-tree analysis.

| Genome | Transcript projections filtered during initial pass | |
|---|---|---|
| | Chimpanzee | Gorilla |
| Outgroup filtering | 43709 | 31678 |
| Best-hit filtering | 13567 | 15765 |

**Table S2:** Number of transcripts filtered out in the initial `pslCDnaFilter` step of CAT, which attempts to remove paralogs and processed pseudogenes.

#### S1.4 Paralogy-filtering evaluation

##### S1.4.1 Alignment of 12 Boreoeutherian genomes

We ran two versions of Cactus (commits `450da74` [best-hit filtering] and `aca859f` [outgroup filtering]) using the following tree:

`(((((Human:0.006969,Chimp:0.009727):0.025291,Rhesus:0.044568):0.07,Tree_shrew:0.19):0.03,(Kangaroo_rat:0.17,(Mouse:0.072818,Rat:0.081244):0.11):0.150342):0.02326,((Dog:0.07,Cat:0.07):0.087381,((Pig:0.06,Cow:0.06):0.104728,Horse:0.05):0.05):0.04);`

Coverage statistics from the resulting alignments were obtained using the `halCoverage` tool.

##### S1.4.2 Annotation using CAT

We produced two alignments using Cactus on the UCSC hg38, panTro6, and gorGor5 assemblies using the same Cactus versions mentioned above. We ran the CAT pipeline at commit `7a8c7e24`, using the GENCODE V30 gene set [4]. We projected the transcripts solely via transMap without the use of the AUGUSTUS modes. Multiple-mapping statistics as well as the gene composition of the final gene set were taken from the `filter_tm_metrics.json` file in the CAT output.

##### S1.4.3 Duplication-timing evaluation

The duplication-timing evaluation was performed using a custom pipeline (`https://github.com/joelarmstrong/treeBuildingEvaluation`) designed to sample columns from a HAL file and eval-

| Genome | Coding genes missing from final set | | Coding transcripts missing from final set | |
|---|---|---|---|---|
| | Outgroup filtering | Best-hit filtering | Outgroup filtering | Best-hit filtering |
| Chimpanzee | 1716 | 1612 | 6244 | 5872 |
| Gorilla | 1829 | 1647 | 6469 | 6100 |

**Table S3:** Number of human genes / transcripts that have no assigned ortholog in the “consensus” CAT gene set across the different alignments.

uate their trees against an independently re-estimated tree of the same region. For this analysis we used the the two 12-boreoeutherian alignments described above, sampling 10,000 columns from the human genome. The comparison trees were built from a context of 1000 bases around the entries in each sampled column using FastTree [13] 2.1.10 and the `-gtr -nt` options. Only duplicated columns were counted in the final output (columns containing no duplications did not count in the results). The coalescence pairs were evaluated using the `--onlySelf` option, meaning that only pairs that included the sampled site were counted in the results. To avoid weighting columns with a high number of copies per genome more than columns with a low number of copies per genome, only a single coalescence was randomly sampled per column.

#### S1.5 Micro-indel events within the 600-way

We extracted all insertion and deletion events by running the `halBranchMutations` tool on every branch in the 600-way alignment. The ungapped insertion and deletion calls (represented by "I" and "D" respectively within the output file) were filtered so that only calls spanning less than 20bp (in the child for insertions, and the parent for deletions) were counted. The rate for each branch was then obtained by dividing the count of these micro-indel events by the total amount of sequence present in the child.

#### S1.6 Generation of the 600-way alignment

The 200 Mammals (200M) alignment was composed of two sets of genomes: newly assembled DISCOVAR assemblies and Genbank assemblies. The DISCOVAR genomes were masked with RepeatMasker [16] commit `2d947604`, using Repbase [1] version 20170127 as the repeat library and CrossMatch as the alignment engine. The pipeline used is available at `https://github.com/joelarmstrong/repeatMaskerPipeline`. The guide-tree topology was taken from the TimeTree database [8], and the branch lengths were estimated using the least-squares-fit mode of PHYLIP [3]. The distance matrix used was largely based on distances from the 4d site trees from the UCSC browser [5]. To add those species not present in the UCSC tree, approximate distances estimated by Mash [9] to the closest UCSC species were added to the distance between the two closest UCSC species. The final guide tree is embedded in the HAL file, and available using the `halStats --tree` command.

The 363 assemblies in the B10K alignment comprised four sets: 236 newly sequenced species for the "family" phase of the project, assembled using SOAPdenovo2 and AllpathsLG, 42 assemblies already sequenced from the "order" phase of the project, 36 assemblies taken from GenBank, and 49 assemblies contributed by other research groups. For the avian guide-tree, we used a tree that the B10K consortium derived as preliminary data from ultraconserved elements.

Both alignments were run on the AWS cloud over the course of 3 weeks for the avians and 2 months for the mammals, using a maximum of 240 `c3.8xlarge` instances and 20 `r3.8xlarge` instances. Because Toil's autoscaling mode was used, this capacity was only fully utilized during the initial phase of the alignment, when the potential for parallelism was at its highest.

The 600-way alignment was formed by aligning the two roots of the B10K and 200M alignments, using the xenTro9 (frog), latCha1 (coelacanth), and danRer11 (zebrafish) assemblies as outgroups. This created a "linker" alignment connecting the roots of the two alignments. The B10K and 200M alignments were then added to this linker alignment using the `halAppendSubtree` command.

#### S1.7 Repetitive elements within ancestral sequences

We ran RepeatMasker [16] on all ancestral assemblies of human within the 600-way alignment (using RepBase [1] version 20170127, selecting the "primate" repeat library and choosing CrossMatch as the alignment engine). We additionally ran the same pipeline against human (as existing annotations used the "Homo_sapiens" repeat library). All ancestors up to human-rhesus had over 78% of the human complement of L1PA6 elements (Figure S3).

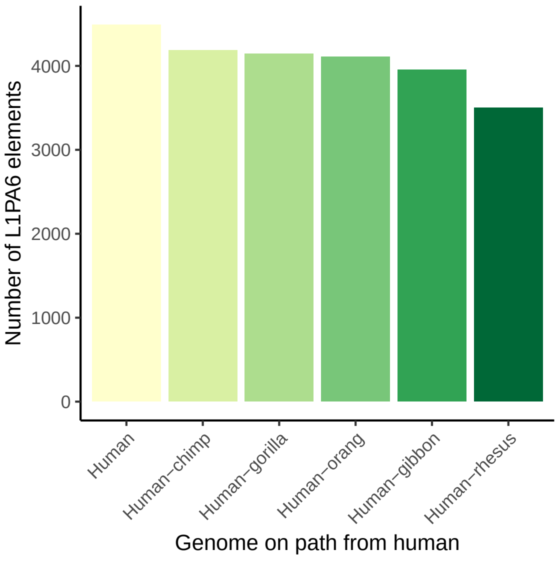


**Figure S3:** Number of L1PA6 elements within ancestral genomes.

#### S1.8 Removing recoverable sequence

The original CAF algorithm described in [12] was focused on removing small rearrangements, while retaining as much of the original alignment relationships as possible in the filtered cactus graph. However, because the input local alignments are insensitive, the original alignment relationships are likely to have missed certain homologies. This can result in what we term *incomplete blocks*: blocks that contain some alignment relationships but are missing others, i.e. are proper subsets of what the corresponding "true" alignment block. In our anchor-and-extend process, once a block becomes an anchor it can never be modified. As a result, these incomplete blocks will remain incomplete: they prevent the true alignment relationship from being found, even if an adjacent syntenic anchor block is complete and contains all desired alignment relationships. These problematic incomplete blocks become more prevalent at longer evolutionary distances: the local aligner will miss more true homologies at increasing distances, causing more incomplete blocks and in turn a far worse alignment.

To remove these incomplete blocks, Cactus originally relied on a heuristic that identified blocks that were "likely" to be incomplete, removing blocks which did not have alignment relationships between all ingroups. However, this heuristic performed poorly in the presence of deletions or missing data: any large deletion in one ingroup could cause huge stretches of the other ingroup(s) to be left unaligned. To remedy this, we have developed a new alteration to the CAF algorithm, one that now focuses on maximizing the potential size of the alignment graph *after* extension as opposed to *before* extension. We call this addition *removing recoverable chains*, because it identifies chains in the cactus graph that represent alignments which could be recovered by the extension process.

The algorithm is applied as a post-processing step after the CAF process described in [12], which proceeds as normal. After the cactus graph is created and filtered, the algorithm identifies *recoverable blocks*. Each block is composed of segments, each of which represent a non-overlapping region of a sequence and which strand is being aligned; we briefly review the necessary terminology, but see [11] for additional context. We call a segment $a$ *left-adjacent* to another segment $b$ if $a$ represents the positive strand and $b$ comes before $a$ in their sequence and there is no other segment between them. Similarly, we call $a$ left-adjacent to $b$ if $a$ is on the negative strand and $a$ comes before $b$ in their sequence ordering with no other intervening segment. If $a$ is left-adjacent to $b$, then $b$ is *right-adjacent* to $a$.

A block is called *recoverable* if, in the case that the block were removed, all its regions would be contained entirely within a single end alignment in the BAR extension phase. The end alignments are identified by looking at all unaligned sequence between the adjacent segments of a single *end* of a block: in short, two end alignments are created for every block, one for all sequence between each segment and its left-adjacent segment, and similarly for the right-adjacent segments. In practice, this means that for some block $A$, it is recoverable if all its segments are all left- or right-adjacent to segments from the same block $B \neq A$.

Whether a block is recoverable depends only on its immediate neighboring blocks. However, it is interesting to consider the maximum set of recoverable blocks, and, by contrast, of unrecoverable blocks — these unrecoverable blocks represent a minimal set of anchors that can be extended from to recover the alignment relationships from the original sequence graph as well as potential additional alignment relationships.

Since the chains and nets within the cactus graph represent a hierarchy of the rearrangements implicit in the alignment, they are helpful for finding this a smaller set of anchors to extend from. We consider what anchors could provide recoverability to a block: if a block $A$'s segments would lie within the end alignment of $B$ if all the recoverable blocks between $B$ and $A$, including $A$, were destroyed, we call *$A$ recoverable given $B$*. The relationship is transitive: if block $A$ is recoverable given block $B$, and $B$ is recoverable given $C$, then $A$ is recoverable given $C$. All blocks in a chain are recoverable given each other, since all blocks in a chain are collinear with each other, potentially with intervening rearrangements located further down the chain/net hierarchy. Similarly, if any block in a chain is recoverable given another block above the chain in the chain/net hierarchy, the entire chain is recoverable given that block. Due to this fact, in order to determine the recoverability status of all blocks, we only have to examine the blocks at the ends of chains and their immediate neighbors, rather than every block.

Though in principle we would need to keep only one block within even unrecoverable chains (since all other blocks within the chain would be recoverable given that single block), to save computational effort in realignment we only destroy or keep entire chains as a unit. In the same spirit, to avoid spending needless effort when the chain is recoverable but very likely is not *incomplete*, we apply a heuristic and do not remove chains that contain the same number of copies in all ingroups and outgroups.

After identifying and removing all recoverable blocks, some blocks previously marked unrecoverable may become recoverable (because adjacent blocks were removed). For this reason, we run the process of identifying and removing recoverable chains multiple times in a loop, until either no recoverable chains are identified or a limit on the number of cycles is reached. The structure of the cactus graph may change after removing recoverable blocks, so we recompute the cactus graph after every removal step. The process that is followed is described in Algorithm S1.

#### S1.9 Improvements from removing recoverable sequence

To quantify the effect that the process of removing recoverable chains (described above) had on real alignments, we ran alignments on a set of 9 Euarchontoglires genomes with the feature turned on and off. The tree used was:

`(((((((human:0.00877,gorilla:0.008964):0.009693,orang:0.01894):0.015511,rhesus:0.037601):0.07392,tarsier:0.1114):0.034014,tree_shrew:0.19114):0.002,(kangaroo_rat:0.171759,(chinese_hamster:0.14,mouse:0.132282):0.11015):0.114051)euarchontoglires:0.020593,(cow:0.18908,dog:0.13303):0.032898);`

We used Cactus commit `56874bde`, with the `--root euarchontoglires` option so that cow and dog were used only as outgroups. Coverage on human increased for all genomes when recoverable chains were removed, especially for those most distant from human (Figure S4). This likely reflects the fact that though the losses caused by not removing recoverable chains in any single subproblem are relatively small, they can compound to be quite significant in large alignments, since many subproblems are involved in creating the alignment between distant species (such as human and mouse, which are separated by 7 internal nodes in this tree).

**Algorithm S1** Recoverable-chain destruction

```
function RemoveRecoverableChains(G, n)
    for 1 ... n do
        cactusGraph ← CreateCactusGraph(G)
        RecoverableChains ← ∅
        for chain C in cactusGraph do
            if
                ▷ A single adjacent end offers the potential for recoverability
                (|C.leftAdjacencies| = 1 or |C.rightAdjacencies| = 1)
                ▷ Shared adjacencies indicate a non-recoverable rearrangement
                and C.leftAdjacencies ∩ C.rightAdjacencies = ∅
                ▷ Links between chain ends indicate a non-recoverable duplication
                and C.leftEnd ∉ C.rightAdjacencies
                    then
                RecoverableChains ← RecoverableChains ∪ {C}
            end if
        end for
        if |RecoverableChains| = 0 then
            break
        else
            Destroy each chain in RecoverableChains
        end if
    end for
end function
```

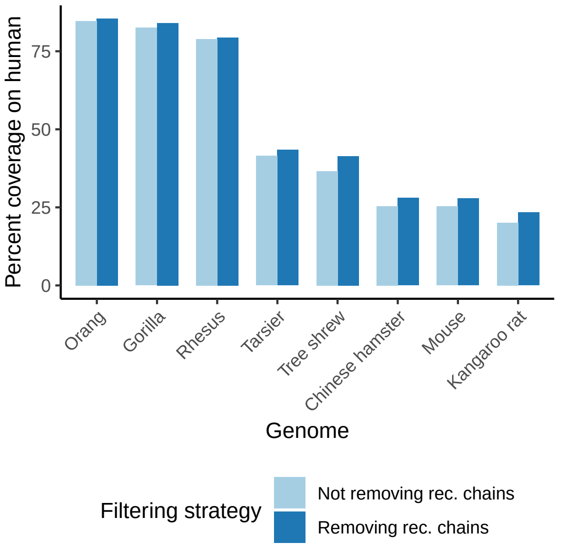


**Figure S4:** Coverage (on the human genome) from alignments with and without removing recoverable chains after the CAF process. While the coverage is increased overall across all genomes when removing recoverable chains, the increase is relatively larger in more distant species.

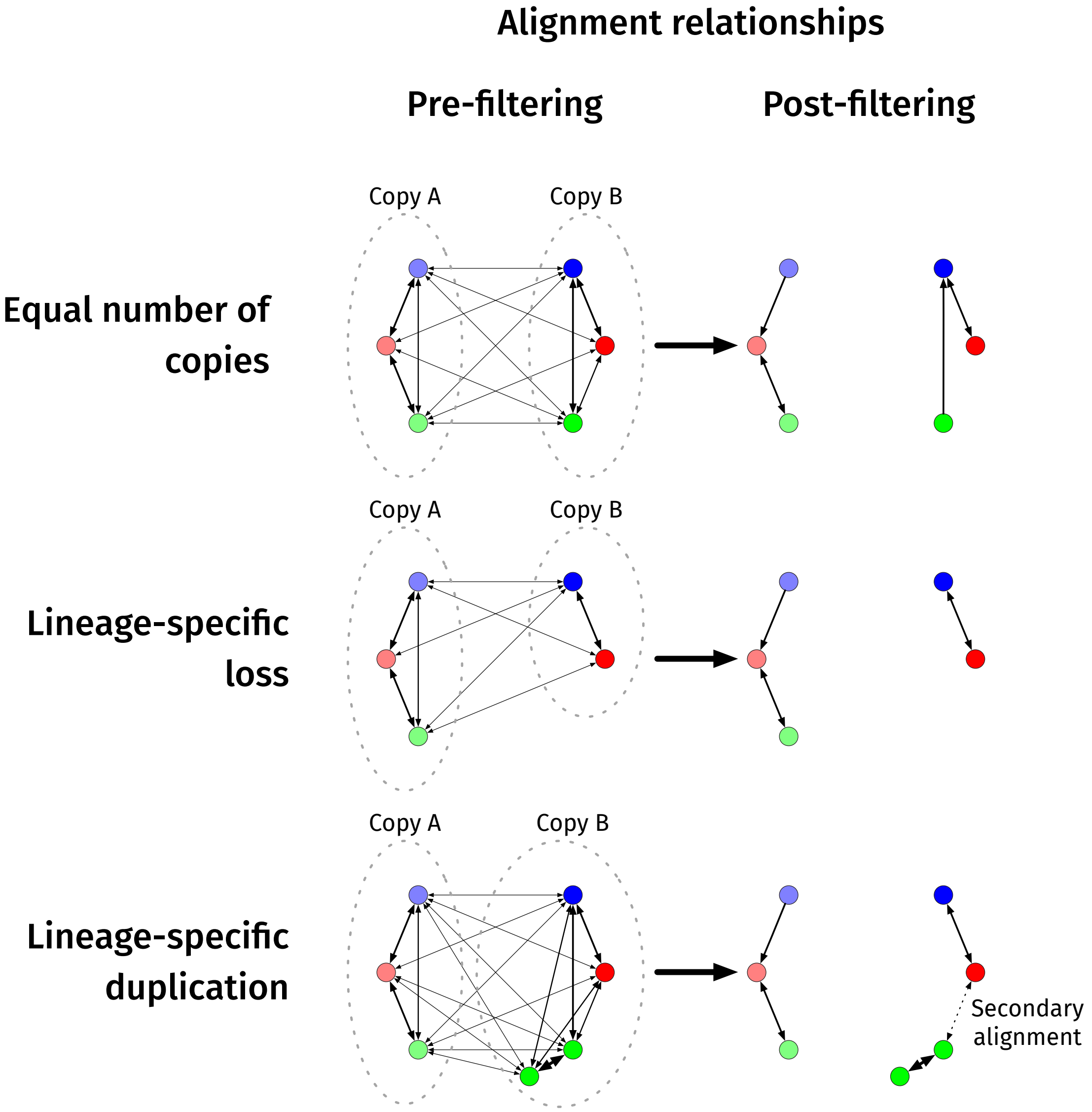


**Figure S5:** A visualization of the best-hit filtering method. Here, each node of the directed graph indicates a single base, and edges represent pairwise alignment relationships (the color of the node indicates the species the base belongs to, and higher thickness of edges represents higher scores of the pairwise alignments). Since Cactus's alignment columns represent the transitive closure of the input pairwise alignment relationships, the final alignment relationships will be represented by connected components within this graph. Taking the single best hit (so that this graph contains at most one outgoing edge per base) results in the correct separation between copies if orthologous copies have higher score, but some lineage-specific duplications require secondary, non-best-hit alignments to bring together orthologs from different species.

### References


[1] W. Bao, K. K. Kojima, and O. Kohany. Repbase Update, a database of repetitive elements in eukaryotic genomes. *Mob DNA*, 6:11, 2015.

[2] R. C. Edgar, G. Asimenos, S. Batzoglou, and A. Sidow. Evolver: a whole-genome sequence evolution simulator. `https://www.drive5.com/evolver`.

[3] Joseph Felsenstein. PHYLIP: Phylogeny Inference Package (Version 3.2). *Cladistics*, 5:164–166, 1989.

[4] Adam Frankish, Mark Diekhans, Anne-Maud Ferreira, Rory Johnson, Irwin Jungreis, Jane Loveland, Jonathan M Mudge, Cristina Sisu, James Wright, Joel Armstrong, et al. Gencode reference annotation for the human and mouse genomes. *Nucleic acids research*, 47(D1):D766–D773, 2018.

[5] M. Haeussler, A. S. Zweig, C. Tyner, M. L. Speir, K. R. Rosenbloom, B. J. Raney, C. M. Lee, B. T. Lee, A. S. Hinrichs, J. N. Gonzalez, D. Gibson, M. Diekhans, H. Clawson, J. Casper, G. P. Barber, D. Haussler, R. M. Kuhn, and W. J. Kent. The UCSC Genome Browser database: 2019 update. *Nucleic Acids Res.*, 47(D1):D853–D858, Jan 2019.

[6] Melissa J. Hubisz, Katherine S. Pollard, and Adam Siepel. PHAST and RPHAST: Phylogenetic analysis with space/time models. *Briefings in Bioinformatics*, 12(1):41–51, 2011.

[7] Erich D Jarvis, Siavash Mirarab, Andre J Aberer, Bo Li, Peter Houde, Cai Li, Simon YW Ho, Brant C Faircloth, Benoit Nabholz, Jason T Howard, et al. Whole-genome analyses resolve early branches in the tree of life of modern birds. *Science*, 346(6215):1320–1331, 2014.

[8] Sudhir Kumar, Glen Stecher, Michael Suleski, and S. Blair Hedges. TimeTree: A Resource for Timelines, Timetrees, and Divergence Times. *Molecular Biology and Evolution*, 34(7):1812–1819, 04 2017.

[9] B. D. Ondov, T. J. Treangen, P. Melsted, A. B. Mallonee, N. H. Bergman, S. Koren, and A. M. Phillippy. Mash: fast genome and metagenome distance estimation using MinHash. *Genome Biol.*, 17(1):132, 06 2016.

[10] Emmanuel Paradis, Julien Claude, and Korbinian Strimmer. APE: Analyses of Phylogenetics and Evolution in R language. *Bioinformatics*, 20(2):289–290, 01 2004.

[11] B. Paten, M. Diekhans, D. Earl, J. S. John, J. Ma, B. Suh, and D. Haussler. Cactus graphs for genome comparisons. *J. Comput. Biol.*, 18(3):469–481, Mar 2011.

[12] Benedict Paten, Dent Earl, Ngan Nguyen, Mark Diekhans, Daniel Zerbino, and David Haussler. Cactus: Algorithms for genome multiple sequence alignment. *Genome research*, 21(9):1512–1528, 2011.

[13] M. N. Price, P. S. Dehal, and A. P. Arkin. FastTree 2–approximately maximum-likelihood trees for large alignments. *PLoS ONE*, 5(3):e9490, Mar 2010.

[14] R. O. Prum, J. S. Berv, A. Dornburg, D. J. Field, J. P. Townsend, E. M. Lemmon, and A. R. Lemmon. A comprehensive phylogeny of birds (Aves) using targeted next-generation DNA sequencing. *Nature*, 526(7574):569–573, Oct 2015.

[15] Liam J. Revell. phytools: an r package for phylogenetic comparative biology (and other things). *Methods in Ecology and Evolution*, 3(2):217–223, 2012.

[16] Smit, A. F. A. and Hubley, R. and Green, P. RepeatMasker Open-4.0. `http://www.repeatmasker.org`, 2013-2015.
